# Ribosomal Biogenesis Hyperactivation and ErbB signalling Mediated Network Rewiring Causes Adaptive Resistance to FGFR2 Inhibition

**DOI:** 10.1101/2024.10.27.620536

**Authors:** Tao Zhang, Sung-Young Shin, Callan McCrimmon, Mandy Theocharous, Ralf B Schittenhelm, Thierry Jarde, Roger J Daly, Lan K Nguyen

**Affiliations:** Department of Biochemistry and Molecular Biology, School of Biomedical Sciences, Monash University, Clayton Victoria 3800, Australia; Biomedicine Discovery Institute, Monash University, Clayton Victoria 3800, Australia; Monash Proteomics and Metabolomics Platform, Monash University, Clayton Victoria 3800, Australia; Department of Anatomy and Developmental Biology, Monash University, Clayton, VIC 3800, Australia; Development and Stem Cells Program, Monash Biomedicine Discovery Institute, Clayton, VIC 3800, Australia; Computational Systems Oncology Program, South Australian immunoGENomics Cancer Institute (SAiGENCI), The University of Adelaide, Adelaide, SA 5005, Australia

**Keywords:** FGFR2 inhibitor, adaptive resistance, proteome, ribosome biogenesis, ErbB/MAPK signalling

## Abstract

Fibroblast Growth Factor Receptor 2 inhibition presents a promising therapeutic approach for restraining the growth and survival of cancer cells, particularly in breast tumours. However, the emergence of resistance to FGFR2 inhibitors like PD173074 highlights the importance of understanding the molecular mechanisms driving resistance and identifying effective therapeutic strategies. In this study, we employed temporal quantitative proteomics and phosphoproteomics, complemented by computational clustering, PTM-SEA analysis, and kinase activity prediction, to monitor the response of MFM223 triple negative breast cancer cells to FGFR2 inhibition. Strikingly, we observed a marked enrichment of ribosome biogenesis function modules immediately following treatment, a phenomenon not observed with other inhibitors. Additionally, we discovered that CX-5461, an RNA polymerase I inhibitor, synergistically enhanced the growth inhibition induced by PD173074, mechanistically attributed to its significant suppression of rDNA transcription stimulated by PD173074. Moreover, our phosphoproteomics dynamic profiling identified the clustering of kinases within MAPK and ErbB signalling pathways, indicative of their reactivation in response to FGFR2 inhibition. Experimentally validating this finding, we observed a notable rebound in phosphorylation levels of key kinases such as ERK1/2 and ErbB3, and demonstrated a substantial synergistic effect of PD173074 in combination with Trametinib, a MEK inhibitor, in suppressing cancer cell growth. Collectively, our findings provide critical insights into the network rewiring triggered by FGFR2 inhibition and offer a foundation for the rational design of combinatorial therapeutic strategies to overcome resistance mechanisms associated with FGFR2 inhibitors.

## Introduction

Despite significant advances in understanding breast cancer’s molecular underpinnings and developing targeted therapies, it continues to be a major contributor to cancer-related morbidity and mortality worldwide ^1–3^. The heterogeneity and complexity of breast cancer underscore the need for a comprehensive understanding of its molecular drivers and mechanisms of therapeutic resistance to develop more effective treatment strategies ^4^.

Among the molecular alterations driving breast cancer progression, dysregulation of the fibroblast growth factor receptor (FGFR) signalling pathway has emerged as a critical factor in a subset of patients ^5,6^. The FGFR family, comprising four transmembrane receptor tyrosine kinases (FGFR1-4), plays pivotal roles in regulating fundamental cellular processes such as proliferation, differentiation, migration, and angiogenesis ^7^. While FGFR aberrations occur in 5-10% of all human cancers, specific cancer types such as urothelial carcinoma and intrahepatic cholangiocarcinoma show an elevated incidence of 10-30% ^5,8^. In breast cancer, FGFR1/2 amplification rates range from 7% to 23% ^9^. These alterations have spurred the development of FGFR-targeted therapies, with several FGFR inhibitors already approved for specific cancer types ^9,10^, such as urothelial carcinoma and cholangiocarcinoma ^10,11^. Yet, their effectiveness in breast cancer is still being investigated. Despite their therapeutic potential, the emergence of drug resistance presents a significant hurdle to the long-term efficacy of FGFR inhibitors ^5,12^.

Resistance to targeted therapies can arise through a variety of mechanisms, including genetic and non-genetic alterations, as well as factors within the tumour microenvironment ^13^. Of particular interest is adaptive resistance, where cancer cells undergo rapid adaptation of their signalling networks to counteract the effects of treatment, often occurring within hours to days of drug exposure. This phenomenon of drug-induced network rewiring has been observed across various targeted therapies and cancer types ^14^. For instance, in BRAF-mutant melanoma, inhibition of BRAF leads to rapid reactivation of the MAPK pathway through CRAF ^15^. In HER2-positive breast cancer, treatment with lapatinib results in the upregulation of FOXO3a and estrogen receptor signalling ^16^. Specifically for FGFR inhibitors, previous studies have reported adaptive resistance mechanisms involving the upregulation of PI3K/AKT pathway ^17–19^, reactivation of MET ^20,21^, and MET-mediated activation of Pyk2 ^22^.

Understanding early adaptive responses is crucial for overcoming resistance to targeted therapies. However, traditional approaches focusing on genetic alterations or static pathway analyses may fail to capture the dynamic and complex nature of adaptive resistance. To address this, unbiased global approaches are needed to elucidate the temporal changes in cellular signalling networks following drug treatment. Global phosphoproteomics profiling offers a powerful tool for capturing these dynamic cellular responses, capable of revealing time-dependent alterations in signalling networks and cellular processes that may contribute to drug resistance. Unlike static studies that may miss transient but crucial adaptive responses, time-resolved phosphoproteomics can provide a comprehensive view of the evolving cellular state after drug treatment.

Despite the clinical promise of FGFR inhibitors, a comprehensive understanding of the global resistance mechanisms to FGFR inhibition in breast cancer is lacking. Moreover, the potential interplay between FGFR inhibition and adaptive signalling changes remains to be fully elucidated. Addressing these knowledge gaps is crucial for developing more effective treatment strategies and overcoming resistance to FGFR-targeted therapies.

In this study, we aim to globally elucidate the molecular mechanisms underlying resistance to FGFR2 inhibition in breast cancer, with a particular focus on early adaptive responses and network rewiring. We employ temporal quantitative proteomics and phosphoproteomics to capture dynamic cellular changes following FGFR inhibition in FGFR2-amplified breast cancer cells. Our findings reveal unexpected links between FGFR inhibition and various cellular processes, providing new insights into the complexity of adaptive resistance mechanisms. Specifically, our findings reveal rapid upregulation of ribosome biogenesis-related genes and hyperactivation of ribosome biogenesis pathways in response to FGFR inhibition. Additionally, we identify significant rebound in ERK, MAPK, and ErbB pathways following FGFR inhibition. We propose that this ribosome biogenesis activation occurs via bypass signalling through the MAPK and ErbB3 pathways. Furthermore, we demonstrate the potential of targeting these adaptive responses through rationale and synergistic drug combinations. Specifically, combining PD173074 with either the ribosome biogenesis inhibitor CX-5461 or the MEK inhibitor Trametinib offers promising strategies to enhance the efficacy of FGFR therapy in breast cancer.

Overall, our study not only advances our understanding of resistance to FGFR-targeted therapies but also provides a framework for investigating adaptive resistance mechanisms in other cancer types and targeted therapies. By unravelling the intricate interplay of signalling pathway rewiring, our work contributes to the broader goal of developing more effective and durable targeted therapies for cancer treatment.

## Results

### Temporal Proteomic and Phosphoproteomic Profiling of FGFR2 Inhibition

Cancer cells often develop adaptive resistance to targeted therapies through rapid network rewiring, a process that occurs without genetic alterations ^14,23–25^. To uncover these dynamic adaptive responses, we employed temporal quantitative proteomic and phosphoproteomic profiling, focusing on the short-term cellular changes following FGFR2 inhibition in breast cancer cells ^26^.

We treated MFM223 cells, a triple-negative breast cancer cell line with FGFR2 amplification, with 20 nM of the FGFR2 inhibitor PD173074. This concentration was chosen based on previous studies reporting IC50 values ranging from 20 nM to 880 nM ^27–33^, and confirmed by our cell viability assays showing ∼50% growth inhibition at 96 hours post-treatment (Figure S1A). Cells were harvested at 0, 1, 4, 9, and 24 hours post-treatment to capture both immediate and adaptive responses (Figure 1A). We then performed total proteome analysis using Liquid Chromatography coupled with Tandem Mass Spectrometry (LC-MS/MS) on 5% of the cell lysates. The remaining 95% was used for phosphoproteome analysis using the EasyPhos method ^34^. This approach allowed us to simultaneously monitor changes in protein expression and phosphorylation status across the proteome.

**Figure 1.**
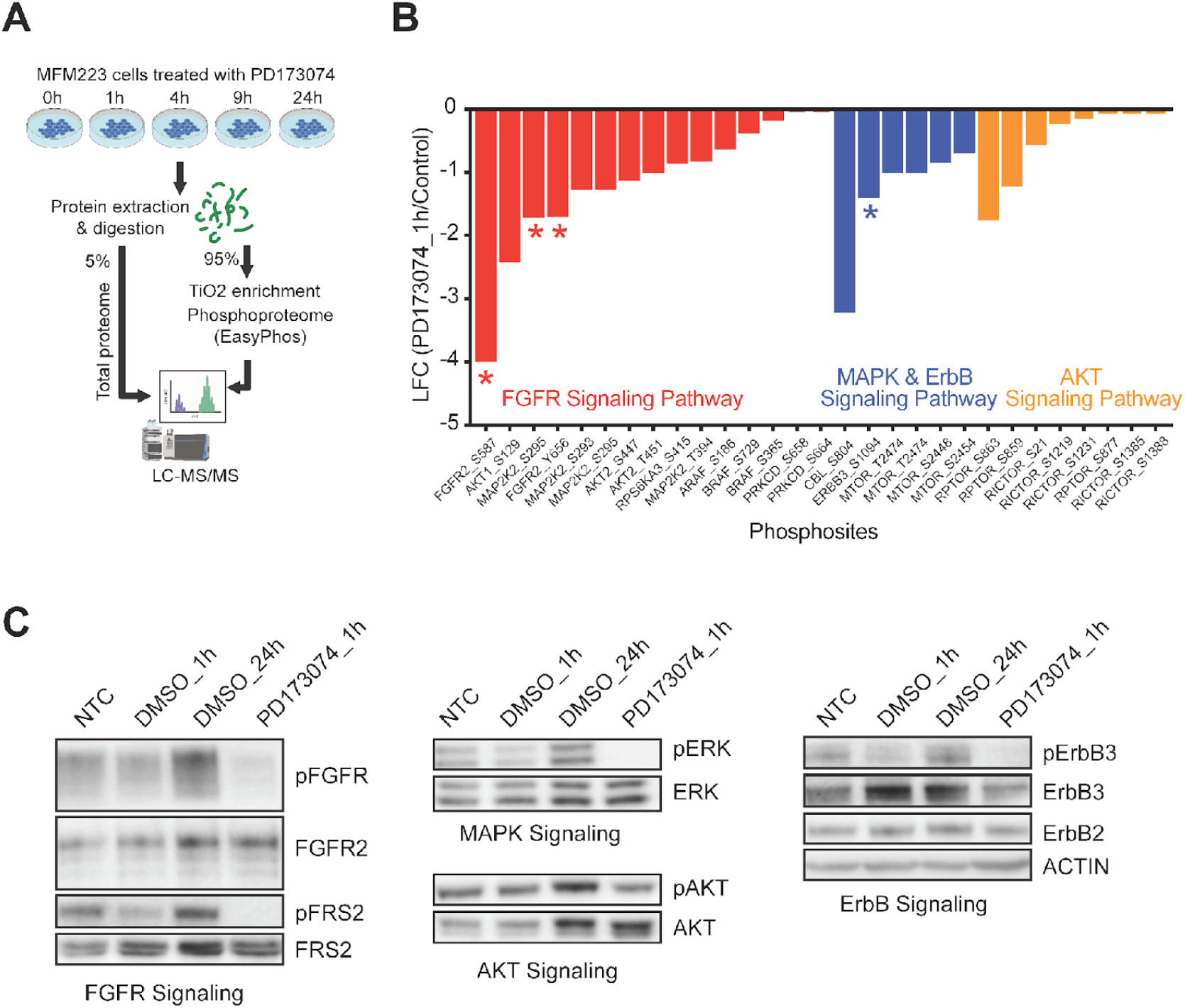
Effect of the FGFR2 inhibitor PD173074 on FGFR downstream signalling pathways. **(A)** Workflow of proteome and phosphorporteome studies. MFM223 cells were treated with 20 nM of PD173074 and harvested at 1, 4, 9, and 24 hours post-treatment in triplicates, with untreated cells as a control. The cell lysates were divided into two parts: 5% for total proteomic profiling and 95% for phosphoproteomic profiling. **(B)** Phosphorylation status of major components of FGFR and its downstream MAPK and AKT signalling pathways was evaluated with Log2 Fold Change (LFC) of phosphosite intensity abundances. * indicates p-value of < 0.05, comparing 1h post-treatment (PD173074_1h) with untreated control (Control). **(C)** Effect of PD173074 treatment. Expression and suppression of FGFR and its downstream signalling proteins at 1 and 24h post-treatment with 20 nM of PD173074, with untreated and DMSO treatment as controls.

To ensure data quality, we conducted rigorous quality control analyses. Principal component analysis and correlation studies of biological replicates demonstrated high reproducibility within time points (r > 0.9 and tight clustering, Figure S1B-D). On the other hand, biological replicates from different time points exhibited distinct separation and minimal correlation (r < 0.1, Figure S1B-D). This analysis confirmed the reliability of our profiling approach ^35^. Consequently, we identified a total of 7,030 proteins and 22,155 phosphosites. After applying statistical filters (p < 0.05, and log2 fold change ≥ |1.5|), we quantified 5,535 proteins and 15,782 phosphosites across all timepoints, with a false discovery rate (FDR) below 1%.

To validate our phosphoproteomic data, we first examined the phosphorylation status of FGFR2, the direct target of PD173074, and its downstream effectors in the AKT and ERK signalling pathways. As expected, we observed significant decreases in the phosphorylation levels of FGFR2 (S587 and Y656), AKT pathway components (AKT-S129, AKT-S447, RPTOR-S863, and RPTOR-S589), and ERK pathway components (MAP2K2-S295 and ErbB3-S1094) at 1-hour post-treatment (Figure 1B). To further confirm these findings, we performed immunoblotting for phosphorylated FGFR, FRS2, ERK, AKT, and ErbB3 in MFM223 cells treated with 20 nM PD173074 for 1 hour (Figure 1C). The immunoblotting results corroborated our phosphoproteomic data, demonstrating effective suppression of FGFR2 and its downstream signalling pathways. Collectively, this comprehensive time-resolved phosphoproteomic profiling approach, coupled with targeted validation, provides a robust foundation for subsequent analyses aimed at uncovering the dynamic cellular responses to FGFR2 inhibition in breast cancer cells.

### Temporal Proteomic Profiling Reveals Ribosome Biogenesis as a Key Adaptive Response to FGFR2 Inhibition

To gain a comprehensive understanding of the cellular response to FGFR2 inhibition, we employed a multi-faceted bioinformatic analysis approach on our temporal proteomic dataset. This integrated strategy combines gene set and pathway enrichment analysis, dynamic pattern classification, and protein interaction network analysis to provide a holistic view of the dynamic changes occurring after drug treatment (Figure 2A).

**Figure 2.**
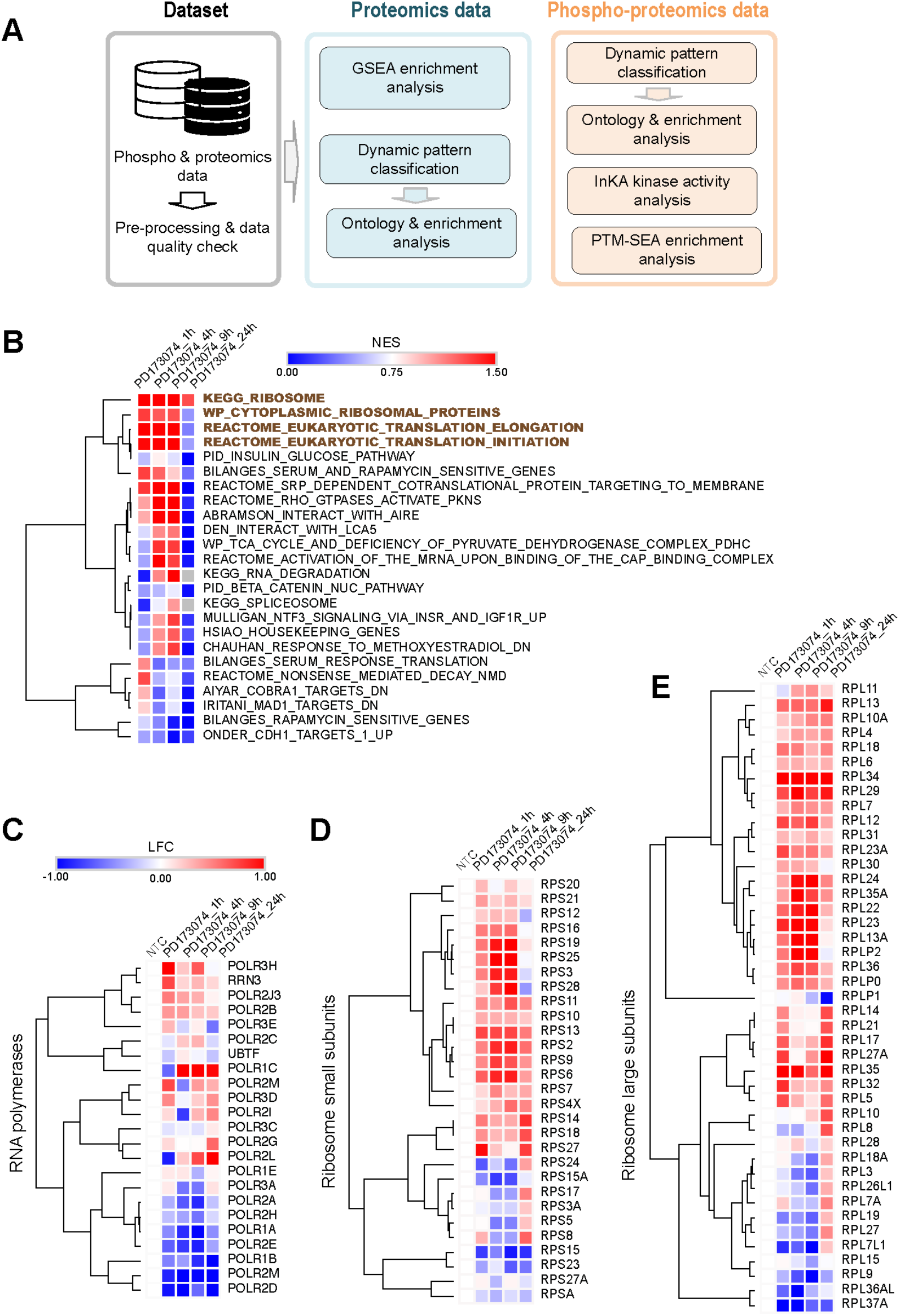
Upregulation of ribosome biogenesis identified by pathway and enrichment analyses. **(A)** Computational workflow and analytical tools employed in this analysis. **(B)** GSEA was performed on dynamic proteomic profiles from MFM223 cells treated with PD173074 for 1h, 4h, 9h and 24h. Results are presented as normalized enrichment scores (NES), with FDR < 0.05. **(C-E)** A heatmap view of LFCs of key RNA polymerase proteins (C), ribosome small subunits proteins (D), and large subunits proteins (E) of dynamic proteomic profiles from MFM223 cells treated with PD173074 at 1, 4, 9, and 24 hours post-treatment, compared to untreated controls (NTC).

We first applied Gene Set Enrichment Analysis (GSEA) on our proteomic dataset (n = 5535 proteins) to identify significantly altered biological pathways over time ^36^. Strikingly, we observed a consistent and significant enrichment of ribosome-related and protein translation-related pathways across all time points following FGFR2 inhibition (Figures 2B, S2A). Specifically, the KEGG ribosome pathway showed persistent upregulation throughout the time course, while pathways involved in translation elongation, initiation, and cytoplasmic ribosomal proteins exhibited upregulation until 9 hours post-treatment, followed by a slight decrease at 24 hours (Figures 2B).

To further elucidate the intricate mechanism underlying ribosome biogenesis, we examined the expression patterns of RNA polymerases I, II, and III, which play complementary and crucial roles in rRNA synthesis and ribosomal protein production (Figure 2C). Note that RNA Polymerase I predominantly synthesizes rRNA, RNA Polymerase II is responsible for producing ribosomal proteins, and RNA Polymerase III transcribes 5S rRNA and tRNAs ^37^. Additionally, we analysed the expression of both small (40S) and large (60S) ribosomal subunit components, which are fundamental for the production of functional ribosomes ^38,39^ (Figure 2D-E). Remarkably, we observed widespread upregulation of these components in response to FGFR2 inhibition, with particularly pronounced effects on both ribosomal subunits.

Next, to characterise the temporal dynamics of protein expression changes more precisely, we employed a dynamic pattern classification approach (Figure 2A). After rigorous data filtering and normalization (Material and Methods, Figure 3A), we clustered differentially expressed (DE) proteins into five distinct groups based on their dynamic profiles: Decrease (DEC, 24.45%), Increase (INC, 10.91%), Down-Then-Up (DTU, 34.59%), Up-Then-Down (UTD, 24.57%) and Fluctuating (FLU, 5.49%) (Figure 3B). Given that the Increase and Down-Then-Up (or rebounding) dynamic patterns of oncoproteins or pro-survival proteins are often associated with drug resistance to the initial treatment ^40^, we focused our subsequent analyses on these groups. Our Gene Ontology (GO) and pathway enrichment analyses of the DTU and INC groups revealed a significant enrichment of ribosome biogenesis-related processes, including ribosomal small subunit biogenesis, rRNA metabolic processes, and RNA processing (Figure 3C), corroborating our GSEA findings (Figure 2B-E). Strikingly, the KEGG ribosome pathway ranked as the most significantly enriched pathway among all identified pathways, reinforcing the central role of ribosome biogenesis in the cellular response to FGFR2 inhibition (Figure S2B).

**Figure 3.**
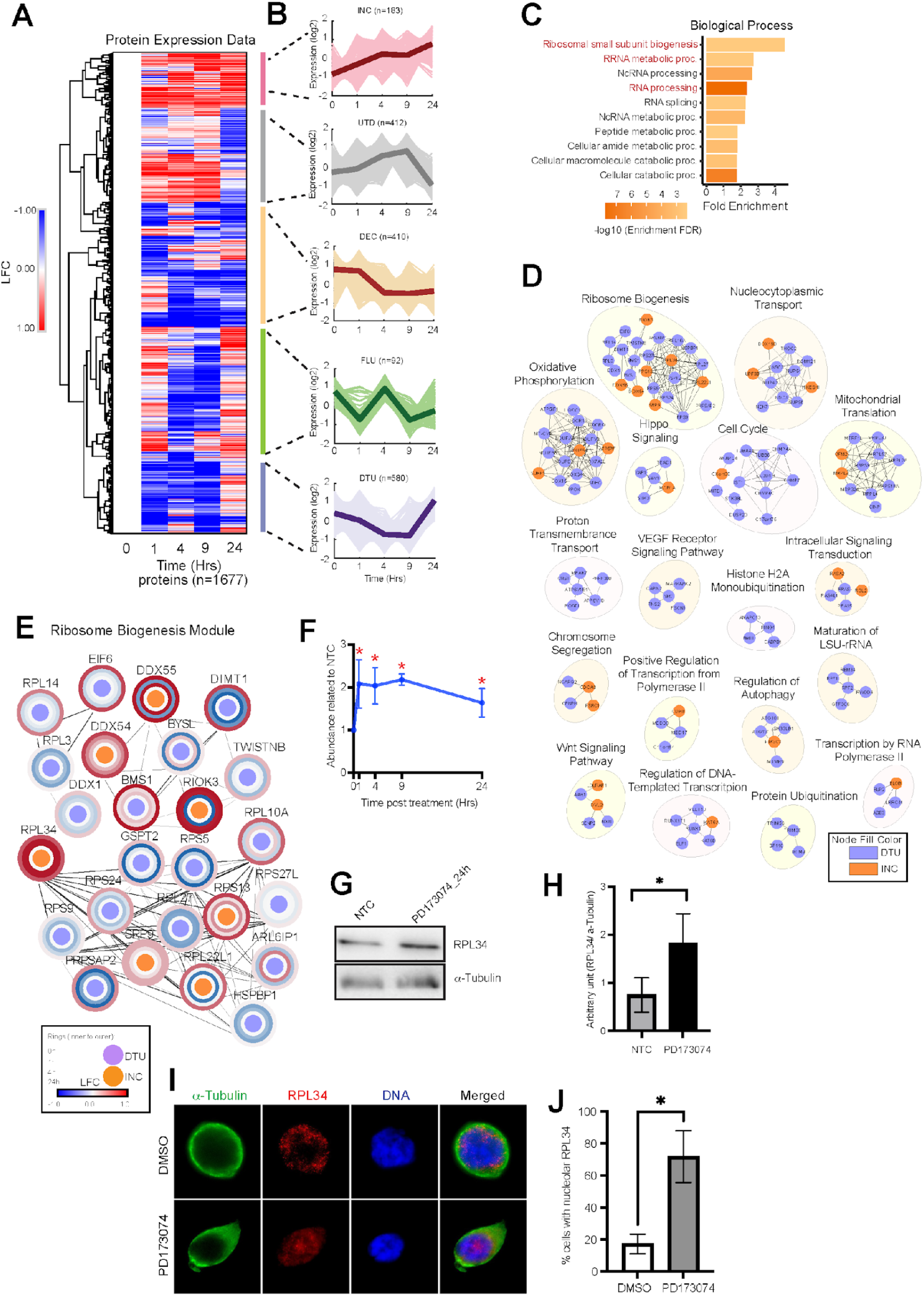
Dynamic pattern classification reveals enrichment of ribosome biogenesis. **(A)** Hierarchical clustering of differentially expressed proteomics profiles. LFC is a log fold change. **(B)** Five dynamic pattern groups. Expression levels are presented in a log2 scale. **(C)** Gene ontology analysis of DTU and INC groups highlighted significant enrichment of ribosome-related biological processes. **(D)** Functional clustering of differentially expressed proteins in DTU (blue) and INC group (orange). **(E)** Detailed analysis of Ribosome Biogenesis Module identified in (D). Rings from inner to outer indicate LFCs of 0, 1, 4, 24 hrs post treatments. **(F)** RPL34 abundance changes over time in response to PD173074 treatment. NTC is untreated control. Error bars mean ± standard error of three biological replicates. * p-value < 0.05. **(G-H)** RPL34 expression of MFM223 cells treated with 20 nM PD173074 for 24h (G) and quantification of immunoblotting of RPL34 with β-Tubulin as a control (H). Representative of three independent experiments. * p-value <0.05. Error bars represent mean ± standard error of three biological replicates. **(I-J)** Immunofluorescence staining of RPL34 and α-Tubulin with DAPI staining for DNA contents. MFM223 cells were treated with 20 nM PD173074 for 4 hrs with DMSO treatment as controls. Representative images of three biological replicates (I). Nucleolar RPL34-positive cells were quantified as a percentage. * p-value < 0.05. Error bars represent mean ± standard error of three independent experiments. At least cells were counted in three independent experiments.

To explore the functional implications of these expression patterns, we constructed a protein interaction network for the aggregated DTU and INC groups using STRING plugin in Cytoscape ^41^ (Material and Methods). Functional enrichment analysis of each protein cluster confirmed that ‘ribosome biogenesis’ is a major functional module, including key proteins such as DDX55, DDX54, BYSL, RPL3, and RPL14 (Figure 3D). A detailed view of this cluster further highlights the rebound and increase patterns of these proteins over time, visualised using Omics Visualizer (Figure 3E). Notably, in addition to functional modules supporting ribosome biogenesis such as ‘maturation of LSU-rRNA’ and ‘nucleocytoplasmic transport’, we also identified other modules including oxidative phosphorylation, Hippo signalling, and cell cycle within these protein clusters, suggesting a complex, multi-faceted cellular response to FGFR2 inhibition (Figure 3D).

To validate our findings in other cellular contexts including both in vitro and in vivo models, as well as with different FGFR inhibitors, we first analysed gene expression profiles from MCF-7 and MDA-MB-231 breast cancer cell lines, RT112 urothelial cancer cell lines, and its PD173074-resistant cells treated with PD173074, obtained from the Connectivity Map platform (CMAP) and public datasets ^42–44^. Consistent with our proteomic data, we observed significant increases in the expression of ribosome large and small subunit genes in these cell lines post-treatment (Figure S2C). In the GO analysis of 1529 up-regulated differential expression genes in PD173074-resistant-RT112 cells, compared with RT112 cells, ribosome biogenesis was significantly enriched (Figure S2D). Similarly, reanalysis of 820 up-regulated differential expression genes in brain tumour glioblastoma PDX mice treated with FGFR inhibitor, Futibatinib, showed that ribosome pathway was significantly enriched (Figure S2E). In contrast, a similar analysis in the MiaPaCa-2 pancreatic cancer cell line treated with a KRAS inhibitor ^45^ did not display any ribosome-related functional modules in the upregulated proteins group (Figure S2F). Taken together, this suggests that the upregulation of ribosome biogenesis is a specific adaptive response to FGFR2 inhibition rather than a general reaction to kinase inhibition.

To further investigate the functional implications of this ribosome biogenesis activation, we focused on RPL34, a ribosomal large subunit protein that showed significant upregulation following FGFR2 inhibition (Figure 3F). Immunoblotting confirmed a substantial increase in RPL34 protein levels 24 hours post-treatment (Figure 3G-H). Moreover, immunofluorescence analysis revealed an increased number of RPL34 foci in the nucleolar regions of treated cells (Figure 3I-J), further supporting enhanced ribosomal subunit synthesis and nucleolar accumulation in response to FGFR2 inhibition.

Collectively, these results revealed a previously unrecognized link between FGFR2 inhibition and enhanced ribosomal biogenesis in MFM223 breast cancer cells. This finding led us to hypothesize that the hyperactivation of ribosome biogenesis may serve as an adaptive response to FGFR2 inhibition, potentially supporting increased protein synthesis demands associated with drug resistance mechanisms ^46^. This hypothesis opens new avenues for understanding and potentially targeting resistance to FGFR inhibitors in breast cancer.

### Targeting Ribosome Biogenesis Overcomes Resistance to FGFR2 Inhibition in Breast Cancer Cells

Our previous findings led us to hypothesize that ribosome biogenesis activation might contribute to the development of resistance to FGFR2-targeted therapies. To explore the therapeutic implications of this discovery, we first assessed the efficacy of PD173074 as a monotherapy. Real-time cell proliferation assays using the xCELLigence Real-Time Cell Analysis (RTCA) system revealed dose-dependent growth inhibition of MFM223 cells by PD173074 over 72 hours, with effects saturating at higher concentrations (Figure 4A). This indicates a potential ceiling effect for PD173074 monotherapy. Furthermore, despite initial growth suppression (Figure 4A), colony formation assays demonstrated the emergence of resistance, as evidenced by persistent colony growth even under PD173074 treatment (Figure 4B-C).

**Figure 4.**
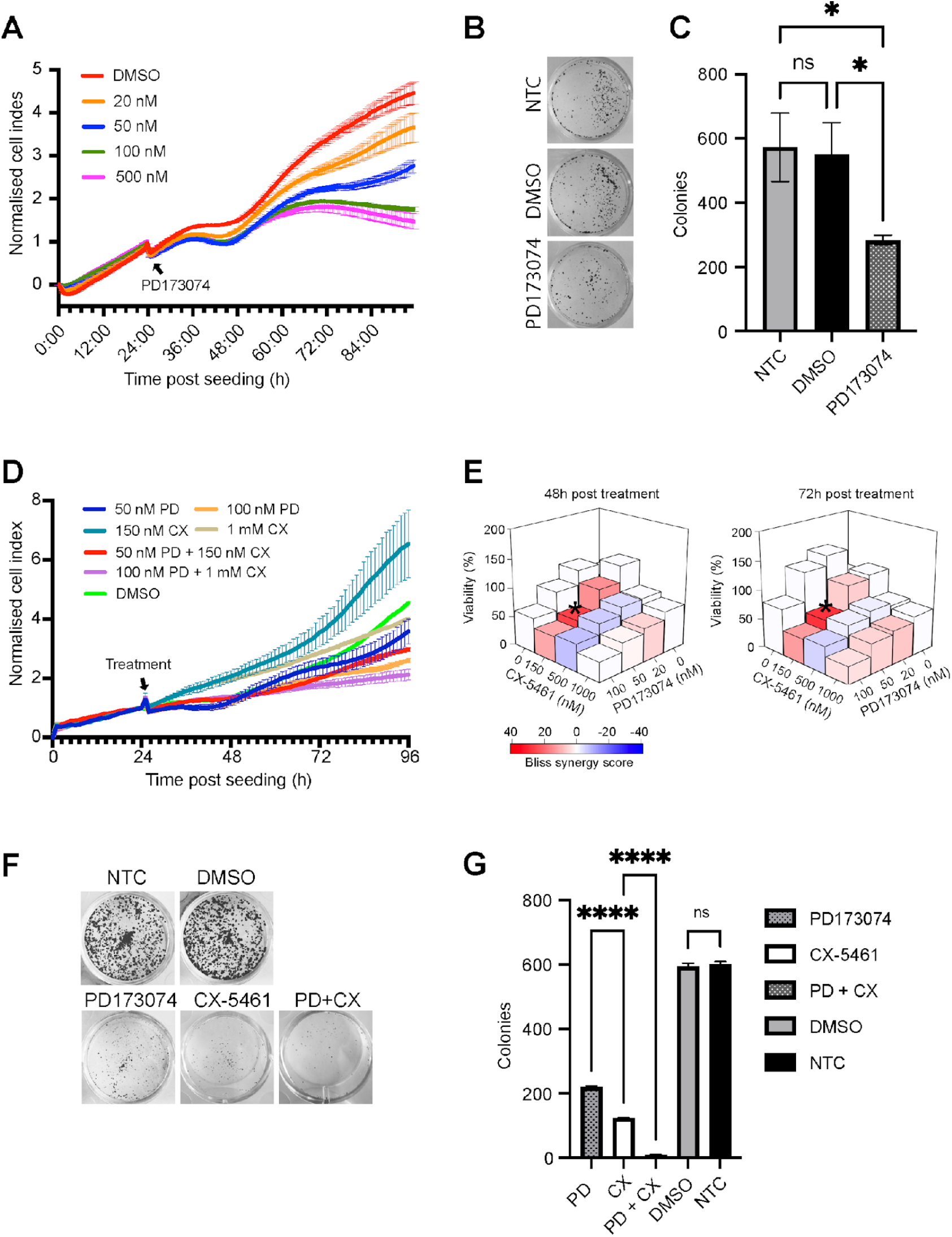
Synergistic effect of PD173074 and CX-5461 on MFM223 cells. **(A)** Measurement of cell proliferation using xCELLigence RTCA system. MFM223 cells were treated with 20 nM, 50 nM, 100 nM and 500 nM PD173074 for 72 hours, with DMSO as controls. Error bars represent mean ± standard error of three independent experiments. Cell index was normalised to the time when the treatment occurred (indicated by an arrow). **(B-C)** Colony formation assays of MFM223 treated with 20 nM PD173074 with NTC and DMSO treatment as controls. Representative images of three independent experiments (B), and colony counts of PD173074 treatment, DMSO treatment, and NTC (C). * p-value < 0.05, ns indicates no significance. Error bars represent mean ± standard error of three independent experiments. **(D)** Cell proliferation measurement using xCELLigence RTCA system for combinational treatment. MFM223 cells were treated with 20 nM, 50 nM, and 100 nM PD173074, and 150 nM, 500 nM, 1 μM CX-5461, and their respective combination for 72 hours, with DMSO treatment as controls. Error bars represent mean ± standard error of three independent experiments. Cell index was normalised to the time when treatment occurred (indicated by an arrow). **(E)** Cell viability and synergy scores quantified for combined treatments of PD173074 and CX-5461 (D), at 48- and 72-hours post-treatment. The synergy score was calculated based on Bliss’ independent model. **(F-G)** Colony formation assays. MFM223 cells were treated with 50 nM PD173074, 150 nM CX-5461, and their combination (PD+CX), with NTC and DMSO treatment as controls. Representative images of three independent experiments (F), and colony counts of PD173074, DMSO, and NTC (G). **** p-value < 0.001, ns indicates no significance. Error bars represent mean ± standard error of three independent experiments.

To evaluate the potential of targeting ribosome biogenesis to overcome this resistance, we assessed the efficacy of combining PD173074 with CX-5461, a potent and selective inhibitor of RNA polymerase I, thereby disrupting rRNA synthesis and ribosome biogenesis ^47^. Indeed, the promising preclinical activity of CX-5461, coupled with its ongoing evaluation in clinical trials across multiple cancers, renders it an attractive candidate for combination therapy ^47–49^. To this end, we employed the xCELLigence RTCA system to monitor cell growth under single-drug and combination treatment conditions. This label-free, impedance-based system allows for real-time, quantitative monitoring of cell proliferation, providing a more comprehensive view of drug effects compared to endpoint assays. Cells were seeded in E-plates and, after 24 hours, exposed to a 4x4 matrix of PD173074 (20-100 nM) and CX-5461 (150 nM-1 µM) concentrations. The combination of PD173074 and CX-5461 significantly reduced cell growth by more than 50% compared to single-agent treatments or DMSO controls (Figure 4D-E). Microscopic examination corroborated these findings, revealing a substantial reduction in cell numbers in the PD+CX treated wells (Figure S3A). To quantify the synergistic effects of these two drugs, we calculated the Bliss Independence score using the SynergyFinder platform ^50^. The most potent synergistic effect was observed at 50 nM PD173074 and 150 nM CX-5461 at 48 and 72 hours post-treatment (indicated by ‘*’, Figure 4E). Interestingly, while CX-5461 has demonstrated potent efficacy with a median IC50 of 38 nM across a panel of cell lines ^47^, it exhibited a relatively reduced potency in MFM223 cells, displaying an IC50 of 501 nM (Figure S3B-C). This observation underscores the importance of combination therapy, as it allows for effective treatment at lower doses of each drug, potentially reducing side effects. To further validate the long-term efficacy of this drug combination, we performed colony formation assays using the optimal doses of 50 nM PD173974 + 150 nM CX-5461 identified in our RTCA experiments. These assays confirmed the sustained effectiveness of the PD173074 and CX-5461 combination in preventing colony formation (Figure 4F-G).

Collectively, these results demonstrate that targeting ribosome biogenesis in combination with FGFR2 inhibition can effectively overcome adaptive resistance, representing a promising combinatorial strategy for improving the efficacy of FGFR2-targeted therapies in breast cancer treatment.

### Differential Regulation of rDNA Transcription by PD173074 and CX-5461 in Cancer Cells

Our previous findings demonstrated a synergistic effect between PD173074 and CX-5461 in inhibiting MFM223 cell growth. To elucidate the molecular mechanisms underlying this synergy, we investigated the impact of these drugs on ribosome biogenesis, specifically focusing on rDNA transcription - a critical initial step in this process ^51^. rDNA transcription produces rRNA within the nucleus, which is subsequently transported to the nucleoplasm and cytoplasm for ribosome maturation. To assess rDNA transcription activity, we utilized two complementary approaches: (i) immunofluorescence analysis of Upstream Binding Factor (UBF) and Fibrillarin, and (ii) direct measurement of nascent RNA synthesis. Note UBF intensity strongly correlates with rDNA transcription activity ^52^, and Fibrillarin, a nucleolar marker, is crucial for DNA transcription and ribosome biogenesis ^53^.

Immunofluorescence analysis revealed that both UBF and Fibrillarin foci intensity increased significantly in PD173074-treated cells compared to DMSO controls (Figure 5A-C), suggesting activation of rDNA transcription. CX-5461 treatment alone significantly reduced both UBF and Fibrillarin foci intensity compared to DMSO controls. Notably, the combination of PD173074 and CX-5461 resulted in a further significant suppression of both markers compared to either drug alone, indicating potent inhibition of rDNA transcription.

**Figure 5.**
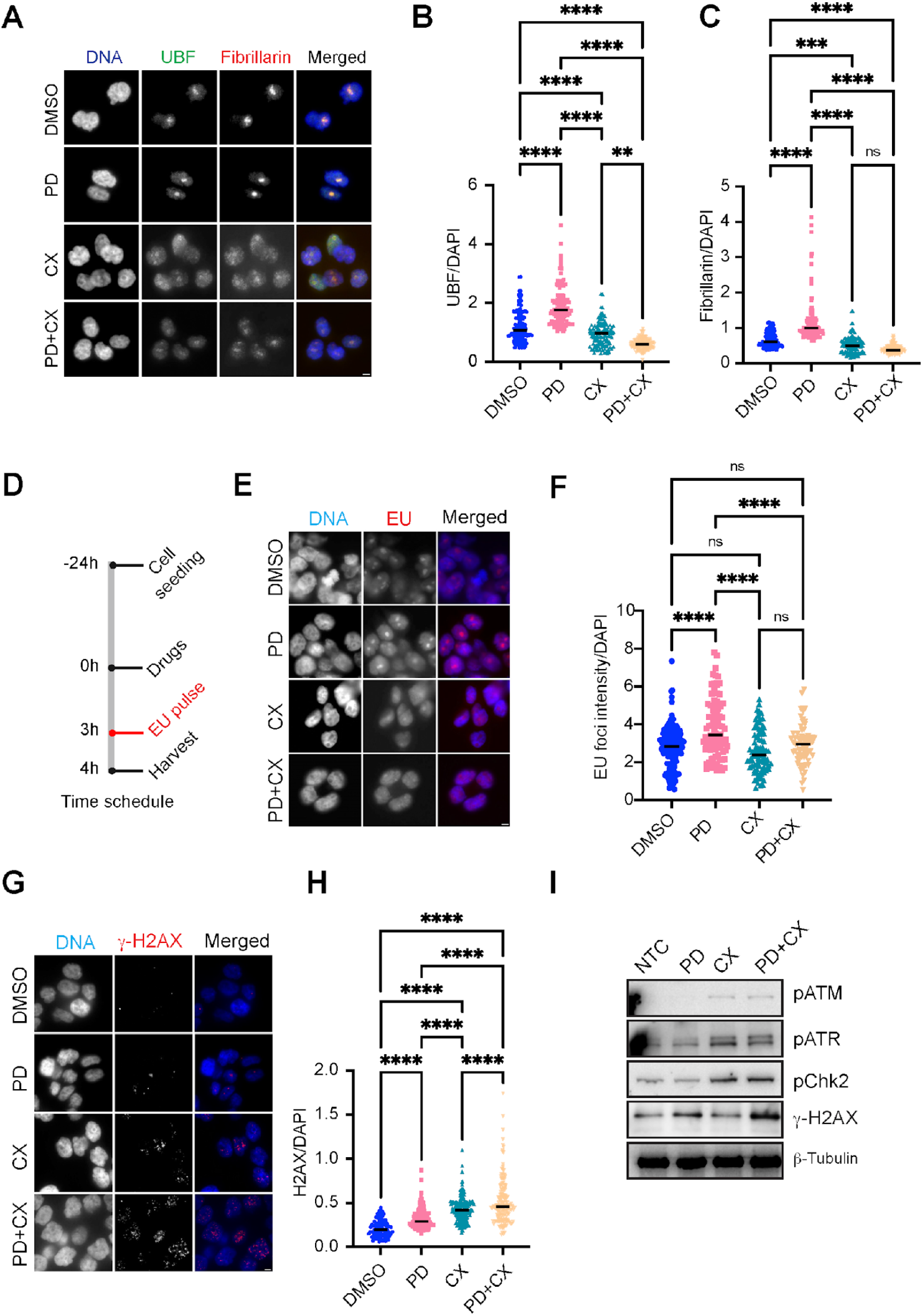
Differential regulation of rDNA transcription and DNA damage checkpoint activation by PD173074 and CX-5461 in MFM223 cells. (A-C) Immunofluorescence staining with UBF and Fibrillarin, with DAPI staining for DNA contents. MFM223 cells were treated with 50 nM PD173074 (PD), 150 nM CX-5461 (CX), and their combination (PD+CX), with DMSO as controls. Cells were harvested at 4 hrs post treatment (A). Representative images of three independent experiments. Quantification of UBF intensity (B) relative to DAPI intensity and Fibrillarin intensity (C) relative to DAPI intensity. **** p-value < 0.001, ** p - value < 0.01, ns indicates no significance. Error bars represent mean ± standard error of three independent experiments. At least 150 cells were measured in each of the three independent experiments. **(D-F)** Workflow of immunofluorescence staining of anti-EU, with DAPI for DNA contents. (D). MFM223 cells were treated with 50 nM PD173074, 150 nM CX-5461 and their combination, with 1-hour pulse of EU and harvested at 4 hrs. Representative images of three independent experiments (E). Quantification of nucleolar EU intensity relative to nuclear DAPI intensity (F). **** p-value < 0.001, and ns indicates no significance. Error bars represent mean ± standard error of three independent experiments. At least 150 cells were measured in each three independent experiments. **(G-H)** Immunofluorescence staining with γ-H2AX, and DAPI staining for DNA contents (G). MFM223 cells were treated with PD173073 and CX-5461 as in (A). Quantification of γ-H2AX intensity relative to DAPI intensity (H). **** p-value < 0.001. Error bars represent mean ± standard error of three independent experiments. At least 150 cells were measured in each of the three independent experiments. **(I)** Western blot analysis for DNA damage responses. MFM223 cells were treated with 50 nM pd173074, 150 nM CX-5461 and their combination (PD+CX), with NTC as controls. Cells were harvested for immunoblotting with pATM, pATR, pChk2, and γ-H2AX, with β-tubulin as loading controls. Representative of two independent experiments.

To more directly quantify rDNA transcription and address potential limitations of the UBF/Fibrillarin assay (such as the presence of UBF foci in mitotic cells), we employed a 5-Ethnyluridine (5-EU) incorporation assay ^54^. This method allows for specific labelling and detection of nascent RNA transcripts, including rRNA in the nucleolus. By thresholding the heterochromatin nucleolus region, we measured 5-EU labelled nascent RNA in the nucleolus as a direct indicator of active rDNA transcription. Consistent with our immunofluorescence data, PD173074 treatment significantly increased nucleolar EU intensity, while the addition of CX-5461 markedly decreased it (Figure 5D-F). These results confirm the ability of CX-5461 to effectively counteract the PD173074-induced upregulation of rDNA transcription in MFM223 cells.

To validate the specificity of PD173074-induced rDNA transcription upregulation, we compared its effects with those of Trametinib, a MEK inhibitor. UBF foci intensity was unaffected by Trametinib treatment but significantly enhanced by PD173074 (Figure S3D), indicating that the observed upregulation of rDNA transcription is a specific response to FGFR2 inhibition rather than a general effect of kinase inhibition.

### Synergistic Induction of DNA Damage and Cellular Stress by Combined PD173074 and CX-5461 Treatment

Given the synergistic inhibition of MFM223 cell growth by PD173074 and CX-5461 and their differential regulation of rDNA transcription, we further investigated the molecular underpinnings of this synergy by examining the effect of these compounds on nuclear structure, DNA integrity, and cellular stress responses.

Immunofluorescence analysis of UBF in pre-extracted cells revealed significant nuclear morphology disruptions in both CX-5461 and combination-treated cells (Material and Methods), characterized by a broken ring-like structure (red arrows, Figure S3E). 3D reconstruction further confirmed distinct nuclear alterations in these conditions, while nucleoli remained intact in PD173074-treated cells (Figure S3F). These observations align with previous reports of CX-5461-induced nuclear structural changes ^55^ and suggest that nucleolar stress may be a primary effect of CX-5461 treatment.

We next hypothesized that these nuclear alterations might lead to DNA damage and activation of cellular stress responses. Indeed, immunofluorescence analysis revealed a profound accumulation of γ-H2AX foci in cells treated with both PD173074 and CX-5461 (Figure 5G). Quantification of the γ-H2AX/DAPI ratio demonstrated a clear stepwise increase in DNA damage from DMSO < PD173074 < CX-5461 < PD+CX, with the combination treatment showing a synergistic effect significantly higher than either drug alone (Figure 5H). Immunoblotting analysis supported these findings, showing dramatically elevated levels of γ-H2AX in combination-treated cells (Figure 5I). Moreover, we observed activation of the DNA damage checkpoint, as evidenced by increased phosphorylation of ATM and ATR in CX-5461 and combination-treated cells, but not in PD173074-treated or untreated cells (Figure 5I).

Collectively, these results suggest that the synergistic anti-proliferation effect of PD173074 and CX-5461 on MFM223 cell is mediated by the induction of profound nuclear stress, extensive DNA damage, and activation of multiple cellular stress response pathways. This combination treatment overwhelms the DNA repair capacity of these cells, resulting in widespread DNA damage and ultimately cell death. This dual mechanism of action, characterized by both growth inhibition and the induction of DNA damage induction, provides a mechanistic rationale for the observed synergy, and suggests that co-targeting FGFR signalling and ribosome biogenesis is an effective strategy to overcome resistance in FGFR2-driven breast cancers.

### Phosphoproteomic Profiling Reveals Dynamic and Diverse Cellular Changes Following FGFR2 Inhibition

To complement our proteomic analysis and gain deeper insights into the dynamic cellular responses to FGFR2 inhibition, we conducted a comprehensive time-resolved phosphoproteomic study. This approach allows us to capture rapid changes in signalling events that may not be reflected in total protein levels, providing a more complete picture of the adaptive responses in MFM223 cells. We analysed 15,782 differentially expressed phosphosites (log2 fold change > |1| compared to untreated controls) across the time course. The majority of phosphosites were on serine residues (87%), with 12% on tyrosine and 1% on threonine (Figure S4A). Upon comparing the DE phosphorylation sites across time points, the majority of phosphosites were regulated more than once (Figure S4B), indicating a dynamic change in the signalling level. Note that certain kinases have phosphorylation at multiple sites, with each site potentially implicated in distinct cellular functions. For example, key regulatory proteins such as TP53BP1, BUD13, and RBBP6 possess five or more sites (Figure S4C), suggesting their potential involvement in multifaceted roles within the cellular response to FGFR2 inhibition. This observation highlights the importance of complementary phosphoproteomic analysis, as this approach provides a depth of functional insight not readily obtainable through proteomic analysis alone.

To gain further insights into the functional implications of these phosphorylation changes, we conducted a series of GO enrichment analyses encompassing (i) Cellular Components, (ii) Molecular Function, and (iii) Biological Process. First, the GO cellular component enrichment analysis found the sustained involvement of key subcellular locations throughout the time course, including the nucleoplasm, nucleolus and cytosol (Figure S5A), highlighting the continuous and interconnected nature of ribosome production ^56,57^. Intriguingly, "Intracellular" terms were significantly enriched only at 1 and 4 hours post-treatment, suggesting rapid activation of intracellular transport pathways. This aligns with the increased demand for protein trafficking associated with enhanced ribosome production ^58,59^. Next, GO molecular function enrichment analysis revealed consistent enrichment of ‘Protein binding’ and ‘Enzyme binding’ functions across all time points, with a notable peak in ‘Nucleic acid binding’ enrichment exclusively at 1-hour post-treatment (Figure S5B). This transient enrichment in nucleic acid binding functions at this early time point aligns with our previous observation of enhanced ribosome biogenesis, as it coincides with the initiation of rDNA transcription, a critical step in ribosome biogenesis (Figure 5). Finally, from the GO biological process analysis we found that enrichment of ‘Ribonucleoprotein complex biogenesis’ (Figure S5C), supporting additional evidence for the sustained activation of ribosome biogenesis pathways following FGFR2 inhibition.

Collectively, these GO enrichment analyses provide a multi-faceted view of the cellular response to FGFR2 inhibition. While strongly supporting our central finding of ribosome biogenesis hyperactivation, they also reveal a complex landscape of cellular adaptations involving transcriptional regulation, intracellular transport, and potential structural remodeling.

### Reactivation of ErbB Signalling Pathways Following FGFR2 Inhibition

Consistent with our prior proteomic analysis, we employed a dynamic pattern classification approach to characterize the temporal dynamics of protein phosphorylation (Figure 6A-B). This analysis identified 2934 differentially expressed (DE) phosphosites across the time course, which were categorized into five distinct kinetic groups: Decrease (DEC, 1.30%), Up then Down (UTD, 86.13%), Down then Up (DTU, 7.57%), Fluctuating (FLU, 2.69%), and Increase (INC, 2.32%) (Figure 6B). As previously discussed, dynamic patterns characterized by Increase and Down-Then-Up (or rebounding) are frequently associated with drug resistance to the initial treatment ^40^. Therefore, we narrowed the focus of our subsequent analyses to the DTU and INC groups.

**Figure 6.**
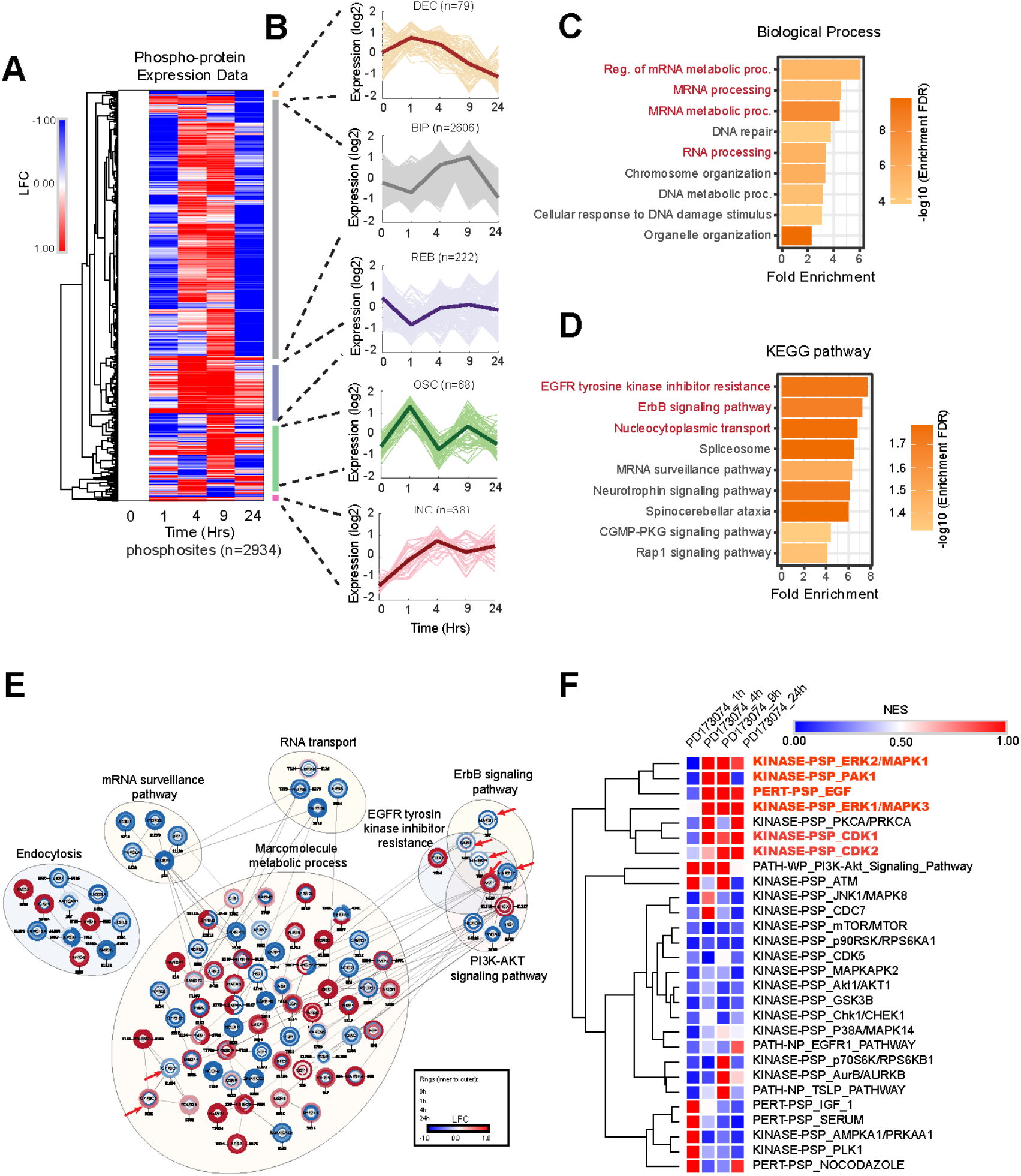
Reactivation of ErbB signalling pathways following FGFR2 inhibition. **(A)** Hierarchical clustering of differentially expressed phosphosites profiles. LFC is a log fold change. **(B)** Five dynamic pattern groups. Expression levels are presented in in a log2 scale. **(C-D)** Gene ontology analysis of DTU and INC groups highlighted significant enrichment of mRNA process and metabolic-related biological processes (C). KEGG pathway enrichment analysis identified EGFR RTK resistance and ErbB associated pathways (D). **(E)** The functional clustering of differentially expressed phosphosites in DTU and INC groups. Rings from inner to outer indicate LFCs of 0, 1, 4, 24 hrs post-treatments. **(F)** PTM-SEA results of dynamic phosphoproteomic profile. NES indicates normalised enrichment score, FDR < 0.05.

To elucidate the functional implications of these dynamic expression patterns, we subjected the DTU and INC dynamic groups to Gene Ontology and pathway enrichment analyses. The GO analysis revealed a significant enrichment of biological processes related to mRNA processing, such as Regulation of mRNA metabolic processing and MRNA metabolic processing, where many components are shared with rRNA metabolic process (Figure 6C). In contrast, the KEGG pathway analysis identified that EGFR tyrosine kinase inhibitor resistance, ErbB signalling pathway, and Nucleocytoplasmic transport pathways are the top 3 enriched pathways (Figure 6D), suggesting a potential reactivation of ErbB signalling pathway.

Next, we assembled a protein interaction network to investigate potential functional relationships. This network was constructed using gene sets identified within the DTU and INC groups. Interaction data were retrieved from the STRING database, and the resulting network was visualized and analysed using Cytoscape software, following the method used in our previous proteome analysis.

Among the 7 major clusters identified, the Macromolecule Metabolic Process cluster was the largest, encompassing 80 proteins. This cluster is primarily implicated in ribosome biogenesis, encompassing the synthesis and assembly of ribosomal proteins and rRNA, the essential macromolecular constituents of ribosomes. Indeed, GTF3C1 and GTF3C2 involved in 5S class rRNA transcription by RNA polymerase III were present in this cluster (red arrows, Figure 6E), which is aligned with our previous results (Figure 3D-E). Additionally, we also found the functional module of ErbB signalling pathway including MAP2K2, MAP2K7, GAB1, EIF4EB1 and AKT1 (Figure 6E), which is consistent with the KEGG pathway enrichment analysis (Figure 6D), suggesting a potential reactivation of ErbB signalling pathways upon FGFR2 inhibition.

To further analyse unique signatures of post-translational modifications we employed Post-Translational Modifications - Signature Enrichment Analysis (PTM-SEA) (Materials and Methods). This analysis identified an enrichment of ErbB/MAPK kinase associated pathways (KINASE-PSP_ERK2/MAPK1, KINASE-PSP_ERK1/MAPK3, KINASE-PSP_PAK1, PERT-PSP_EGF) and cell cycle associated mechanisms (KINASE-PSP_CDK1, and KINASE-PSP_CDK2). These pathways exhibited a pattern of acute inhibition followed by a significant rebound (Figure 6F), supporting a reactivation of ErbB and cell cycle signalling pathways following FGFR2 inhibition.

Inferring kinase activity is essential for understanding the function and adaptation of signalling pathways, particularly in response to perturbations like drug treatment. This knowledge can be leveraged to uncover mechanisms of signal rewiring and design effective therapeutic strategies to overcome resistance to kinase inhibitors. To this end, we employed the Integrative Inferred Kinase Activity ^60^ score algorithm, a robust tool that integrates both kinase-centric analyses, including Kinome and activation loop phosphorylation data, and substrate-centric analyses, such as PhosphoSitePlus and NetworKIN, to rank kinase activity and visualize kinase-substrate networks ^61^. Specifically, we obtained kinase activity scores using the INKA algorithm, which were subjected to clustering and functional annotation within Cytoscape, employing the MCODE and STRING apps to assess and represent the degree of interconnectivity ^62^. Consistent with our analysis in pattern classification and PTM-SEA, we identified ErbB and MAPK signalling pathways as two main clusters, with hyperactive MAPK1/3 (kinase scores) in the ErbB group, and FYN and AKT3 in the MAPK group (Figure 7A). The kinase-substrate networks further supported the reactivation of major kinases such as AKT1, MAPK1, MAPK3, and GSK3B, connected to their known substrates (Figure S6A).

**Figure 7.**
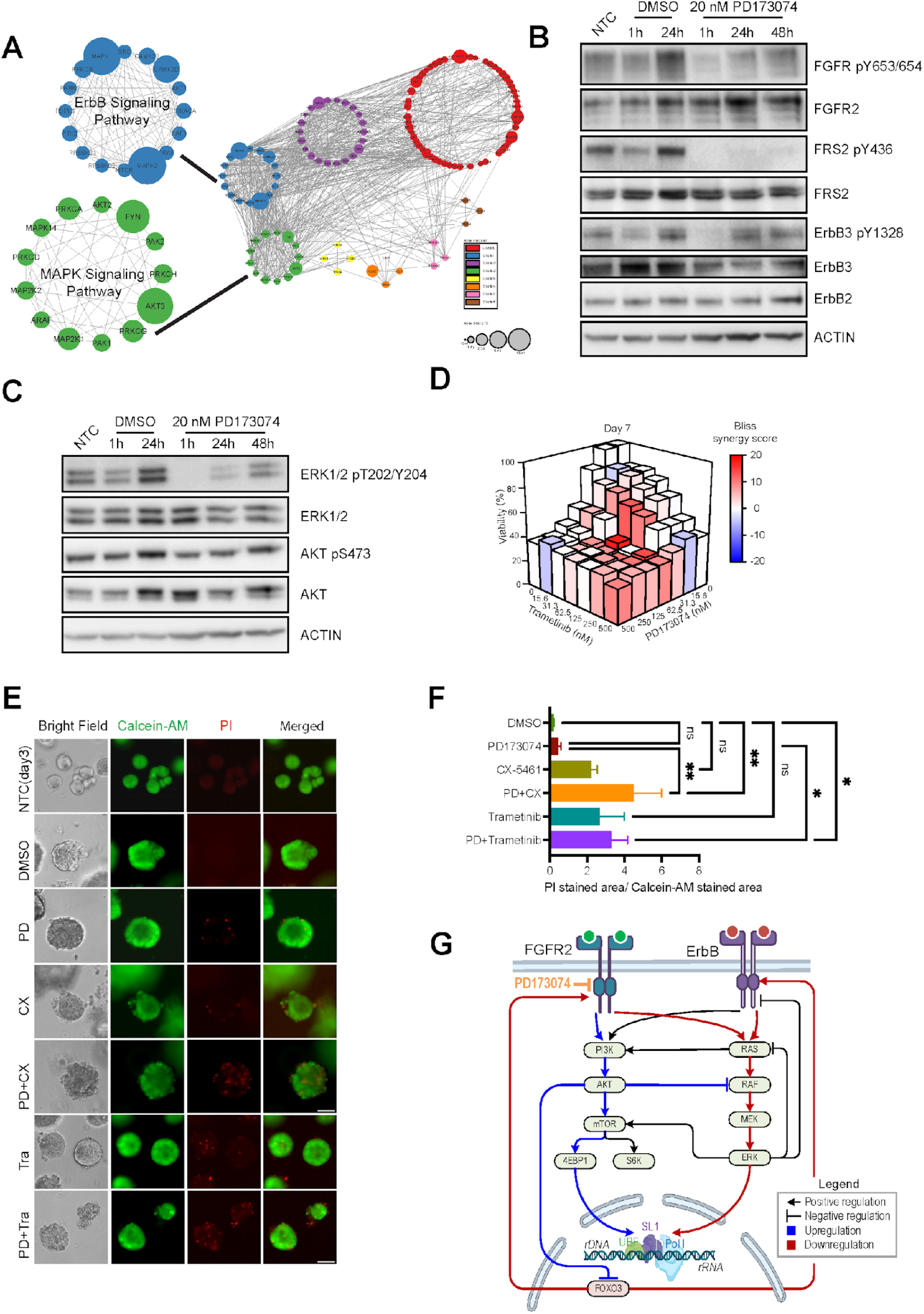
Synergistic effect of PD173074 and Trametinib on MFM223 cells. **(A)** Kinase activity analysis through clustering of InKA score. MAPK signalling and ErbB signalling pathways were highlighted as two major kinases clusters. The node size indicates the LFC of kinases activity of InKA scores at 24 hours post-treatment relative to NTC. **(B-C)** Western blot analysis for RTK and MAPK signalling proteins. MFM223 cells were treated with 20 nM PD173074 at 1 h, 24 h and 48 h post-treatment, with NTC and DMOS treatment as controls. Representative of three independent experiments. **(D)** Cell viability and synergy scores quantified for combined treatments of PD173074 and Trametinib. Cell viability from MTS assays were determined at 7 days post-treatment of PD173074 (0 - 500 nM) and Trametinib (0 - 500 nM). The synergy score was calculated based on Bliss’ independent model. **(E-F)** Effect of combinational treatment on TNBC patient-derived organoids. The organoids were seeded 3 days prior to treatment with 50 nM PD173074, 500 nM CX-5461, 30 nM Trametinib, and their combinations, with NTC (days 3) and DMSO as controls. After 10 days, the organoids were stained with Calcein-AM (live cells, green fluorescence) and propidium iodide (PI; dead cells, red fluorescence) (E). The radio of PI – positive to Calcien-AM-positive area was quantified (F), ** p-value < 0.01, * p-value < 0.05, and ns indicates no significance. Error bars represent mean ± standard error of three independent experiments. **(G)** Resistance to FGFR2 inhibition is mediated by reactivation of MAPK and ErbB signalling pathways, leading to hyperactivation of the ribosome biogenesis pathway.

To experimentally verify the reactivation of ErbB and MAPK signalling pathways, we treated MFM223 cells with 20 nM PD173074, and measured their responses at 1h, 24h, and 48h post-treatment, with DMSO-treated cells as controls. Phosphorylation of ErbB3, ERK1/2, and AKT, key kinases in the ErbB and MAPK signalling pathways, rapidly decreased within 1 hour post-treatment, but recovered after 24h post-treatment, persisting until 48 hours (Figure 7B-C). Importantly, the reactivation of ErbB and MAPK pathways occurred despite the continuous inhibition of FGFR and FRS2 phosphorylation, confirming that the reactivation did not occur due to the drug’s wearing off effect, but rather through a network rewiring. To alleviate concerns about the loss of effectiveness of FGFR2 inhibition following prolonged treatment, we treated cells with 20 nM PD173074 for 24 hours, and the supernatant was collected and used to treat fresh cells again for another 24 hours (called new 24 hours). As expected, FGFR2 inhibition was still effective, as determined by the complete inhibition of FGFR, FRS2 and ERK1/2 phosphorylation after additional 24 hours treatment (Figure S6B).

### Targeting ErbB/MAPK Overcomes Resistance to FGFR2 Inhibition in Breast Cancer Cells

PTM-SEA and INKA analysis indicated that the ErbB/MAPK pathway is reactivated following FGFR2 inhibition, and pERK rebound was experimentally confirmed. Thus, to assess the therapeutic potential of targeting the MAPK pathway in conjunction with FGFR inhibition, we evaluated the impact of combining PD173074 with MEK1/2 specific inhibitor, Trametinib, in MFM223 cells. Trametinib was FDA-approved in 2013 as a monotherapy for patients with metastatic melanoma carrying the V600E mutation. A combination of Trametinib with Dabrafenib (a BRAF inhibitor) has also received FDA approval for the treatment of melanoma with BRAF mutations ^63^. MFM223 cells were treated with a 7x7 matrix of varying concentrations of PD173074 and Trametinib (ranging from 0 to 500 nM) for 4 and 7 days, followed by MTS assays. Combined treatment resulted in a dose-dependent decrease in cell viability (Figures 7D, S6C). To quantify synergism between two drugs, the Bliss Independence model was employed using the SynergyFinder platform ^50^. The combination of 50 nM PD173074 and 100 nM Trametinib exhibited the most pronounced synergistic effect at day 7 (Figure 7D). To further validate the cellular response at the signalling network level, we treated the cells with 50 nM PD173074 and/or 100 nM Trametinib, with untreated cells and DMSO-treated cells served as controls. Cells were harvested at 1h, 24h, and 48h following treatment. Immunoblotting confirmed that this combination effectively inhibited the phosphorylation of ErbB3, inhibiting the rebound of ErbB/MAPK signalling pathways due to FGFR2 inhibition (Figure S6D).

### Evaluation of Drug Combinations in Breast Cancer Patients-Derived Organoids

To visually ascertain the cytotoxic effects of these drug combinations, we employed a patient-derived organoid system. Human TNBC patient-derived organoids were seeded 3 days prior to treatment. The organoids were treated with 50 nM PD173074, 500 nM CX-5461, 30 nM Trametinib, and their respective combination, with DMSO serving as controls. After 10 days treatment, the organoids were stained with Calcein-AM and Propidium iodide (PI) to distinguish live cells (Calcein-AM, green fluorescence) from dead cells (PI, red fluorescence). By 3 days post-seeding, the untreated (NTC) organoids exhibited smooth, intact morphology with no PI staining (Figure 7E). Similarly, DMSO-treated-organoids retained structural integrity with minimal dead cells (PI) staining. However, treatment with the combination of PD173074 and CX-5461 disrupted the organoid structure, significantly increasing the ratio of PI-stained (dead) to Calcein-AM-stained (live) areas, and markedly reducing the sizes of live (Calcein-AM stained) organoid compared to DMSO and PD173074 alone (Figures 7E-F, S7). A similar effect was observed with the combination of PD173074 and Trametinib (Figures 7E-F, S7). These findings collectively indicate that co-targeting ErbB/MAPK or ribosome biogenesis and FGFR2 synergistically enhances cytotoxic effects, overcoming resistance to FGFR2 inhibition in breast cancer cells.

## Discussion

The high frequency of FGFR alterations in various cancers ^5,6^, coupled with FGFR’s important roles in developmental and physiological processes and oncogenesis ^7^, suggest its potential as a promising therapeutic target. However, the multifaceted nature of resistance mechanisms to FGFR inhibitors limits their clinical application ^17–22^. Therefore, a comprehensive understanding of network rewiring in response to FGFR inhibition will facilitate the development of more effective therapeutic strategies. To address this issue, this study utilized temporal quantitative proteomics and phosphoproteomics, integrated with comprehensive bioinformatic analysis and experimental validation, and elucidated that treatment with the RNA polymerase I inhibitor CX-5461 potentiated PD173074-induced growth inhibition. Dynamic phosphoproteomics profiling revealed reactivation of the MAPK and ErbB signalling pathways upon FGFR2 inhibition, followed by experimental validation of ERK1/2 and ErbB3 rebound. Notably, combining PD173074 with the MEK inhibitor Trametinib yielded significant synergistic suppression of cancer cell growth.

Our methodology employed an integrated bioinformatic pipelines, ensuring robust and cross-validated identification of key signalling pathways and molecular processes involved in therapeutic resistance. Specifically, we utilized temporal quantitative proteomics and phosphoproteomics to capture dynamic changes in protein expression and phosphorylation status in FGFR2-overexpressing MFM223 cells treated with FGFR inhibitor PD173074. We then conducted GSEA to reveal the significantly enriched pathways (Figures 2, 3) and post-translational modifications associated with FGFR2 inhibition (Figures 6, 7). Subsequently, these datasets were processed using computational clustering algorithms to identify distinct dynamic patterns of protein and phosphoprotein alterations over time (Figures 3A, 6A). This was complemented by kinase activity prediction to pinpoint active kinases potentially driving resistance mechanisms (Figures 7A, S6A). Our integrative bioinformatic approach, combining high-throughput experimental data with robust computational analyses, provided a comprehensive and cross-validated landscape of the molecular underpinnings of FGFR2 inhibitor resistance.

Ribosome biogenesis is a highly coordinated process involving the transcription, processing, and assembly of ribosomal RNA (rRNA) and proteins ^64^. Dysregulation of biogenesis has emerged as a hallmark of cancer cells, supporting their increased demand for protein synthesis to sustain rapid proliferation and survival ^65^. Targeting ribosome biogenesis using inhibitors such as CX-5461 has shown promise in preclinical models of cancer by inhibiting cancer cell growth and sensitizing cells to chemotherapy ^47^. Indeed, this finding prompted phase I clinical trials (NCT02719977 and NCT04890613), as a fast-track by the United States Food and Drug Administration for Breast and Ovarian cancers with BRCA1, BRCA2, PALB2, or other HRD mutations ^48,49^. Furthermore, a recent study revealed that ribosome biogenesis inhibition triggers hyperactivation of FGFR signalling in glioma cells ^66^. On the other hand, CX-5461 has been shown to stabilize FGFR1 G-quadruplex structures, and diminish FGFR1 expression, thereby limiting cellular plasticity of FGFR1 overexpressing metastatic breast cancer ^67^. However, the precise role of ribosome biogenesis in mediating the cellular response to FGFR inhibition in breast cancer remains to be elucidated.

In our study, we revealed PD173074 treatment in MFM223 cells resulted in increased expression of ribosome large and small subunits, along with upregulation of RNA polymerases (Figures 2B-E, S2A-E), suggesting the hyperactivation of ribosome biogenesis and enhanced transcriptional activity in response to FGFR2 inhibition. These findings were further supported by the change of Upstream Binding Factor (UBF) foci intensity (Figures 5A-B, 5D-F, S3D) and the localization of UBF to nucleolar regions and its association with Fibrillarin, a nucleolar marker (Figure 5C). Interestingly, the reduction in UBF foci intensity upon treatment with CX-5461, an inhibitor of RNA polymerase I, corroborates the specificity of our observations (Figures S3D-E), and implicates UBF as a potential therapeutic target for modulating ribosome biogenesis in breast cancer ^68^. Indeed, targeting rRNA methyltransferase Fibrillarin, another key regulator of rDNA transcription, has shown great potential to overcome chemoresistance by inhibiting ribosome biogenesis ^69^.

An increasing body of evidence suggests that mutation and/deregulation of ribosomal proteins are closely linked to genetic disease such as ribosomopathies, and cancer ^46^. Notably, overexpression of RPL34 has been demonstrated to drive the development and progression of non-small cell lung cancer (NSCLC) cells ^70^, while knockdown of RPL34 is associated with the suppression of the proliferation of gastric cancer cells ^71^. Moreover, RPL34 mutation or dysregulation is also implicated in the development and progression of cancers such as Esophageal, NSCLC, gastric, pancreatic, glioma cells, osteosarcoma and cervical cancers ^46^. Our proteome analysis revealed upregulation of RPL34 in response to FGFR2 inhibition (Figures 2B-E, 3A-F). This finding was validated through immunoblotting and immunofluorescence assays (Figure 3G-J). The observed localization of RPL34 in both nucleoplasm and nuclei suggests an enhanced ribosome biogenesis following FGFR inhibition. Clinically, these results raise the intriguing possibility that RPL34 could be a potential biomarker for hyperactivation of ribosome biogenesis in breast cancer. Supporting this notion, survival analysis using data from The Cancer Genome Atlas (TCGA) Breast Invasive Carcinoma (PanCancer Atlas) demonstrated that high mRNA expression of RPL34 in the patients was significantly correlated with a poorer overall survival rate (HR=0.62, p < 0.05) (Figure S8).

Our phosphoproteome analysis revealed enrichment of mRNA associated biological processes (Figure 6C) and reactivated MAPK and ErbB signalling pathways (Figures 6D-F, 7A), consistent with the observed hyperactivation of ribosome biogenesis pathways identified in our proteomic study (Figures 2B-E, 3A-F). This finding suggest a coordinated signalling network that governs cellular adaptation to FGFR2 inhibition. These results raise the question of how the reactivation of MAPK and ErbB signalling is mechanistically linked to the hyperactivation of ribosome biogenesis. Previous studies provide clues to this connection. Notably, PI3K/AKT and MAPK pathways are known to linked to enhanced ribosome biogenesis ^46^. Thus, the reactivation of MAPK and ErbB pathways represent a compensatory mechanism employed by cancer cells to overcome the inhibitory effects of PD173074 on FGFR2 signalling ^17,18,72–74^. Intriguingly, our data displayed a reciprocal inhibitory relationship between the PI3K/AKT and ERK pathways. Specifically, treatment with the PI3K/AKT inhibitors MK2206 and BYL-719 led to increased pERK, a key protein in the MAPK pathway (Figure S9A-B). Conversely, inhibition of MEK by Trametinib resulted in increased pAKT (Figure S9C). The crosstalk and feedback regulation between PI3K/AKT and ERK pathways have been extensively characterized. AKT phosphorylates and inhibits RAF, thereby attenuating the RAF-MEK-ERK cascade ^75,76^. RAS directly activates PI3K through interaction with its amino-terminal Ras-binding domain (RBD) ^77^. The transcription factor FOXO1/3a contributes to this regulatory network by inducing the expression of RTKs, including FGFR2 and ErbB ^78^. AKT-mediated phosphorylation of FOXO leads to its cytoplasmic sequestration and inactivation ^79^. Consequently, inhibition of AKT allows FOXO nuclear translocation, resulting in the transcription of RTKs. This interplay forms a negative feedback loop ^80^. Indeed, consistent with this feedback regulatory mechanism, our PD173074 treatment displayed increased expression of FGFR2 and ErbB2/3 (Figures 7B, S9D).

The FGFR2 signalling network illustrates that the drug response of PD173074 is modulated by a complex interplay of multiple feedback and crosstalk mechanisms (Figure 7G). PD173074-mediated FGFR2 blockade rapidly suppresses PI3K/AKT and RAS activity. However, this concurrently relieves AKT-mediated negative regulation of RAF, leading to MEK/ERK pathway activation. Simultaneously, the release of FOXO3 inhibition contributes to the transcriptional upregulation of ErbB and FGFR2, thereby activating a feedback loop; notably, ErbB enhances RAS/RAF/MEK/ERK signalling. Consequently, ERK-driven ribosome biogenesis constitutes a major resistance mechanism to FGFR2 inhibition (Figure S10). On the other hand, treatment with the MEK inhibitor, Trametinib, directly inhibits the ERK pathway, leading to suppressed ribosomal biogenesis activation via inhibition of both ERK and mTOR (Figure S10). Note that ERK partially activates mTOR ^81^. This mechanism explains the synergistic effect observed in our experiments upon combining FGFR2 and MEK inhibition (Figure 7D).

While this study provided novel insights into FGFR2 inhibitor resistance mechanism and suggested potential therapeutic strategies to overcome it, limitations warrant consideration. While Erdafitinib and Pemigatinib are newer, potent, and selective inhibitors of FGFR1/2/3, demonstrating significant suppression of cell viability (IC50 ∼ 10 nM at 96h) compared to DMSO-treated controls (Figure S11A-B), and minoring the increase of FGFR2, pAKT rebound and RPL34 upregulation observed with PD173074 treatment (Figure S11C-D), several factors motivated the selection of PD173074 as the FGFR2 inhibitor for this study. PD173074 represented one of the most extensively characterized, widely utilized, and potent FGFR2 inhibitors available ^9,10^ and, thus the extensive body of literature and validated assays associated with PD173074 facilitated leveraging existing knowledge and methodologies ^27,29,82^, ensuring the reliability and reproducibility of our findings. The use of only MFM223 cells in this study might limit the comprehensive assessment of the heterogeneity of responses to FGFR inhibition across different breast cancer subtypes. However, the inclusion of TNBC organoids provide a more physiologically relevant, patient-derived model, thereby increasing the translational relevance of our findings. Future studies incorporating a panel of additional cell lines will be necessary to validate and extend these findings to a broader range of breast cancer contexts.

In conclusion, our findings delineate a complex interplay between FGFR2 inhibition and ribosome biogenesis in breast cancer cells, highlighting the intricate nature of targeted therapy responses. These results provide novel insights into the molecular mechanisms underlying therapeutic resistance and may inform the development of new therapeutic strategies targeting ribosome biogenesis pathways to overcome resistance to FGFR inhibition in breast cancer.

## Methods

### Cell culture and chemicals

MFM223 cells were cultured in RPMI medium, supplemented with 10% (v/v) fetal bovine Serum, 10 μg/mL Actrapid penfill insulin (Clifford Hallam Healthcare), and 20 nM HEPES (Life Technologies), as described previously ^83^. The cells were maintained in a humidified atmosphere incubator at 37 °C with 5% CO_2_. The chemical inhibitors PD173074 (Catology No. HY-10321), CX-5461 (Catology No. HY-13323), and Trametinib (Catology No. HY-10999) were purchased from MedChemExpress through Focus Bioscience.

### Organoid passaging

TNBC organoids (HBC14) were obtained from the Monash BDI organoid program. Organoids passaging was conducted as described before ^84^. Briefly, organoids embedded in Matrigel were mechanically scraped, spun down, and subjected to trypsinization using TrypLE Express (Thermo Fisher Scientific). Cells were centrifuged at 1500 rpm for 3 minutes at 4 °C. After discarding the supernatant, the cell pellet was resuspended in Matrigel, and 50 μl of the cell suspension was seeded in a dome shape in a 24-well plate. Following 10 minutes of polymerisation, complete medium was added on top of the Matrigel. The organoids were maintained in a humidified atmosphere incubator at 37 °C with 5% CO_2_, the media (without Y-27632) were changed every 3 days after passaging.

### Immunoblotting

Immunoblotting was performed as described previously ^85^. Briefly, cell pellets were resuspended in RIPA buffer containing a cocktail of protease inhibitors (Roche), 40 mM β-glycerophosphate, 0.1 mM Na-orthovanadate, and 1mM NaF for 30 minutes on ice. After centrifugation at 140,000 x *g* for 15 minutes at 4 °C, the protein concentration was determined using the BCA method (Pierce 23225). 20 μg protein from each sample were loaded onto 7% or 12% SDS-PAGE gels. Primary antibodies used were pFGFR (Y653/654) (1:1000, CST 3471), FGFR2 (1:500, SCBT sc-6930), pFRS2α (Y436) (1:1000, CST 3861), FRS2 (1:1000, Millipore 05-502), pERK1/2 (Thr202/Tyr204) (1:10000, CST 4370), ERK1/2 (1:1000, CST 4695), pAKT (S473) (1:1000, CST 4058C), pan AKT (1:1000, CST 4685BC), pErbB3 (Y1328) (1:1000, CST 8017S), ErbB3 (1:1000, CST 4754S), RPL34 (1:500, Sigma HPA035139), pATM (Ser1981) (1:1000, CST 5883), pATR (Ser428) (1:1000, CST 2853), pChk2 (Thr68) (1:10000, CST 2197), pHistone H2A.X (Ser139) (1:1000, CST 9781), α-tubulin (1:1000, Sigma T5168) and β-actin (1:2000, SCBT sc-69879). Secondary antibodies used were goat anti-ribbit IgG (H+L)-HRP conjugate (1:3000, Bio-Rad 1707615) and goat anti-mouse IgG (H+L)-HRP conjugate (1:3000, Bio-Rad 1706516).

### Immunofluorescence, EU labelling, microscopy and image analysis

Immunofluorescence staining was performed as described previously ^85^. In brief, cells grown on cover slides were fixed with 4% paraformaldehyde (PFA) for 15 minutes, permeabilised with 0.3% Triton-X100 and blocked with 3% BSA in PBS. For pre-extraction method, cells were pre-extracted with 0.5% Triton-X100 before fixation with 4% PFA. Cells were then stained with primary antibodies including RPL34 (1:400, Sigma HPA035139), γ-H2A.X (phospho S139) (1:800, Abcam), UBF (F-9) (1:400, SCBT sc-13125), UBFT (1:400, Sigma HPA006385), Fibrillarin (1:400, CST 2639T) and α-tubulin (1:800, Sigma T5168). Secondary antibodies used were anti-rabbit Alex Fluor 594 (1:1000, Invitrogen A-11012), and anti-mouse Alex Fluor 488 (1:1000, Invitrogen A32722). The cells were mounted with Vectashield mounting medium containing 10 μg/mL DAPI (Vector Laboratories, H-1000-10) prior to imaging.

To detect nascent RNA by 5-ethynyl uridine (EU) labelling, cells were pulsed with 1 mM EU for 1h before harvest. EU was detected with Click-iT RNA Alex Fluor 594 imaging kits (Invitrogen C10330) following the manufacturer’s procedure.

Immunofluorescent images were captured using a Lecia DMi8 and Lecia Thunder inverted light microscope with a 63x objective. For 3D reconstruction, 45 section images (0.2 μm per section) with a 100x objective were taken and analysed using IMARIS 8.1.2. Fluorescence intensity was measure by ImageJ. Three independent experiments were performed with at least 150 cells per experiments measured. The arbitrary intensity units of Alex 594 or Alex 488 to DAPI were plotted. To determine the nascent rRNA intensity, the fluorescence intensity of rRNA was measured based on nucleolus regions threshold with the DAPI channel ^86^.

### Fluorescence staining of organoids, imaging and analysis

Calcein-AM/PI staining of organoids was conducted as described previously ^87^. After 10 days of treatment, the organoids were stained with Calcein-AM (1:1000, Thermo Fisher Scientific C3099) and Propidium Iodide (1:1000, Thermo Fisher Scientific P3566) for 15 minutes. Imaging was performed the EVOS M5000 imaging system (Thermo Fisher Scientific AMF50000) following the manufacturer’s description. Fluorescent images captured in the green (Calcein-AM) and red (PI) channels were further analysed using Cellprofiler pipelines. Three independent experiments were performed with at least three images captured per sample.

### Colony formation assay

The colony formation assay was performed as described previously ^85^. In brief, 100 cells per well were seeded in triplicate on 12-well-plates. Cells were treated with 50 nM PD173074, and/or 150 nM CX-5461 for 14 days, with untreated and DMSO-treated cells as controls. The cells were replaced with media with/without fresh drugs every 3 days. After 14 days, the plates were placed on ice and washed 2 times with cold PBS. The colonies were fixed with ice-cold methanol for 10 minutes and stained with crystal violet staining solution for 5 minutes at room temperature (RT). The colonies were then washed 3 times with MQ H2O before drying at RT overnight. The colonies were counted as described by ImageJ as described before ^85^.

### MTS assays and Real-Time Cell Analysis

MTS assays were performed as described previously ^83^. In brief, 2000 cells per wells in triplicates were seeded in a 96 well plate overnight. The cells were treated in a 7x7 drug matrix including PD173074 at the concentration of 0, 15.6nM, 31.3nM, 62.5nM, 125nM, 250nM, 500nM and Trametinib at the concentration of 0, 15.6nM, 31.3nM, 62.5nM, 125nM, 250nM, 500nM for 4 and 7 days. Absorbance was measured at the wavelength of 490 nM using Pherastar plate reader. The readings were normalised to DMSO-treated controls and the viability was further subjected into SynergyFinder (https://synergyfinder.org/) for Bliss synergy score calculation ^50^.

Real-Time Cell Analysis was conducted in xCELLigence RTCA system (Agilent) according to the manufacturer’s procedure. In brief, 15,000 cells per well were seeded in duplicate on E-plate 16 PET (Agilent). After 1-hour incubation, this plate was mounted onto xCELLigence system. The impedance in each well was monitored every hour for 24 hours (T=24). The cells were then treated with a 4x4 drug matrix including PD173074 (at the concentrations of 0, 20 nM, 50 nM, and 100 nM) and CX-5461 (at the concentrations of 0, 150 nM, 500 nM and 1 μM) and monitored every hour for another 120 hours. The real time Cell Index was calculated using a built-in algorithm in the xCELLigence system and the Cell Index normalised to the readings at T=24 were plotted over time. Similar to the MTS assays, the viability normalised to DMSO-treated controls was subjected to SynergyFinder for Bliss synergy scores calculation.

### Mass spectrometry-based proteomics

Mass spectrometry-based proteomics were performed as described before ^34,88^. In brief, MFM223 cells were seeded in triplicate in 15 cm petri-dishes overnight, followed by treatment with 20 nM PD173074 for 1, 4, 9 and 24 hours, with untreated cells as controls. After 2 times washed with ice-cold TBS, cells were harvested using freshly prepared SDC lysis buffer (4% sodium deoxycholate, 100 nM Tris-HCl pH 8.5), followed by heat treatment at 95 °C for 5 minutes. After sonication, the protein concentration of lysate was determined by the BCA method. The Reduction/Alkylation buffer (containing 100 mM Tris(2-carboxyethyl) phosphine and 400 mM 2-chloroacetamide pH 7.0) was used to reduce disulphide bonds and carbamidomethylate cysteine residues, followed by heating at 95 °C for 10 minutes. After cooling, digestion with Trypsin and LysC (2 µg of each enzyme per 200 µg of proteins) was setup overnight at 37 °C with shaking at 1,500 rpm. The SDC was precipitated from the solution using 400 µL of isopropanol (ISO), and 100 µL of EP enrichment buffer (containing 48% trifluoroacetic acid and 8 mM KH2PO4) was added to each sample, which was then centrifuged to clear the supernatants (2000 × g, 15 min). The samples were then split for Total Proteome and Phosphoproteome analysis.

### Total proteomic sample preparation

For total proteome analysis, the samples were subjected to a desalting process using C18 spin columns as previously describe (Ref). Subsequently, the samples were dried with a SpeedVac and reconstituted in 2% (v/v) acetonitrile ^89^ and 0.1% formic acid, targeting a peptide concentration of 0.1 μg/μL.

### Phosphoproteomic sample preparation

For phosphoproteome analysis, the supernatants were transferred into fresh Eppendorf tubes after centrifugation, and the enrichment of Phosphopeptides were carried out as described before ^34,88^. In brief, peptides were enriched at a ratio of 12:1 TiO_2_ bead: protein (5010-21315, GL Sciences, Tokyo, Japan) for 5 minutes at 40 °C with shaking (2,000 rpm). The phosphopeptides were eluted with EP elution buffer (5% (v/v) NH4OH in 32% (v/v) ACN) followed by desalted using SDB-RPS (Empore™, CDS Analyticl, Oxford, PA, USA) stage tips prepared in-house and eluted with 20 µl of 25% (v/v) NH4OH in 60% (v/v) ACN. The peptides were then dried using a SpeedVac and resuspended in 2% (v/v) ACN /0.3% (v/v) TFA.

### Mass spectrometry analysis

The UltiMate 3000 RSLC nano LC system (Thermo Fisher Scientific) coupled to a Q Exactive HF mass spectrometer (Thermo Fisher Scientific) was used to analyze the samples. Peptides were loaded onto an Acclaim PepMap 100 trap column (100 µm × 2 cm, nanoViper, C18, 5 µm, 100 Å, Thermo Fisher Scientific) and separated on an Acclaim PepMap RSLC analytical column (75 µm × 50 cm, nanoViper, C18, 2 µm, 100 Å, Thermo Fisher Scientific). For each LC–MS/MS analysis, 1 µg of peptides, as measured by a nanodrop 1000 spectrophotometer (Thermo Fisher Scientific), was loaded onto the pre-column with microliter pickup. Peptides were eluted using a 2-hour linear gradient of 80% (v/v) ACN/0.1% FA at a flow rate of 250 nL/min using a mobile phase gradient of 2.5–42.5% (v/v) ACN. An HRM DIA method was employed, consisting of a survey scan (MS1) at 35,000 resolution (automatic gain control target 5e6 and maximum injection time of 120 ms) from 400 to 1220 m/z, followed by tandem MS/MS scans (MS2) through 19 overlapping DIA windows increasing from 30 to 222 Da. MS/MS scans were acquired at 35,000 resolution (automatic gain control target 3e6 and auto for injection time) using a stepped collision energy of 22.5, 25, and 27.5% and a 30 m/z isolation window. The spectra were recorded in profile type. MaxQuant Version 1.6.0.16 analysis software was used for peptide identification with default settings and an FDR of 1% at peptide level.

### Functional annotation analysis (GSEA, PTM-SEA, INKA, Cytoscape)

Gene Ontology analysis including GO cellular component, GO Biological process, and GO molecular function, were performed using ShinyGO 0.80 ^90^ and the STRING Enrichment app within Cytoscape 3.9.0 ^91^ and plotted with GraphPad Prism 9.0.1. Functionally organised networks were analysed, clustered and visualised using ClueGO and CluePedia ^92^ or EnrichmentMap apps ^93^ covering Gene Ontology, KEGG, WikiPathways and Reactome within Cytoscape 3.9.0. Functional modules were clustered and visualised using the ClusterViz ^94^, ClusterMaker2 ^95^, Omics Visualizer ^96^ apps within Cytoscape 3.9.0. Integrative Inferred Kinase Activity analysis was performed using the online platform (https://inkascore.org) ^61^. Gene sets enrichment with statistically significant difference in samples (1h vs NTC, 4h vs NTC, 9h vs NTC and 24h vs NTC) were determined by Gene Set Enrichment Analysis (GSEA) version 4.2.1 ^36,97^. Post-Translational Modifications - Signature Enrichment Analysis (PTM-SEA) was conducted through the public server GenePattern (https://cloud.genepattern.org/gp/pages/index.jsf) with a plug-in module PTM-SEA ^98^. Heatmaps were generated using an online tool Morpheus (https://software.broadinstitute.org/morpheus/). Venn diagrams were generated using an online tool InteractivVenn (http://www.interactivenn.net) ^99^. Bar charts, pie charts and scatter plots were generated using GraphPad Prism 9.0.1. Three-dimensional plots were generated using plot3D R package (https://cran.r-project.org/web/packages/plot3D/index.html).

### Pattern classification

For dynamic pattern classification of proteins/phosphosites, we first screened for proteins whose expression was statistically upregulated or downregulated at any time point after FGFR2 inhibitor treatment. Then, the time profile data for these proteins was subjected to a min-max normalization, scaling features (proteins) to a range of 0 to 1 using the following formulation:

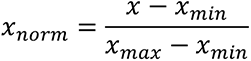

where *x* is the original value. *x_min_* is the minimum value in the dataset for that feature. *x_max_* is the maximum value. *x_norm_* is the normalized value. Thus, the minimum value becomes 0 and the maximum value becomes 1.

The dynamics patterns were clustered into predefined five different groups based on their dynamic profiles using Hierarchical Clustering Algorithm: Up then Down (UTD) – an initial increase followed by a decrease, Down then Up (DTU) – an initial decrease followed by an increase, Fluctuating (FLU) – an initial increase followed by decrease, then increase again, or vice versa, Increase (INC), and Decrease (DEC). For the implementation of this algorithm, the scaled profiles were clustered into fifteen groups using an unsupervised clustering method with MATLAB (2024a) function *cluster* which creates agglomerative clusters from *linkage* function. Then, five predefined labels (UTD, DTU, FLU, INC and DEC) were assigned to each group. Note that we employed the fifteen clusters in our unsupervised clustering as this facilitated straightforward visual assessment of the dynamic patterns. Varying the number of clusters, however, is not anticipated to substantially alter the results.

### Data mining

The signatures of MCF-7 and MDA-MB-231 cells treated with 100 nM and 500 nM PD173074 for 6h and 24h were retrieved from ConnectivityMap (CMAP) (https://clue.io/command?q=/home) ^42^, clustered and visualised with Morpheus, where only ribosomal gene sets were displayed. Expression profiles of RT112 cell and its FGFR inhibitor resistant derivative (GSE201395) were retrieved and reanalysed ^43^. Expression profiles of PDX G39 treated with Futibatinib and DMSO (GSE245624) were retrieved and reanalysed ^44^.

### Survival analysis

Survival analysis utilized mRNA expression and associated clinical data from 1083 breast invasive carcinoma patients, obtained from the TCGA, PanCancer Atlas ^89^ (cBioportal; https://www.cbioportal.org/). Patient samples were stratified into high (> 70% percentile) and low (< 30% percentile) RPL34 expression groups. 324 met criteria for each group (test vs control). The log-rank test was applied to compare overall survival between these groups (p<0.05 considered statistically significant). Kaplan-Meier estimates and survival curves were generated using a web-based application ^100^ (https://can-sat.erc.monash.edu/).

### Statistical analyses

Statistical analyses were conducted using Prism 9.0.1 and included unpaired t-tests or one-way ANOVA. *p* < 0.05 was considered statistically significant.

## Contributions

Conceptualization: T.Z., S.Y.S., and L.K.N. Experimental design and methodology: T.Z., S.Y.S., R.D. and L.K.N. Experiments: T.Z., C.M, and M.T. Analysis: T.Z., and S.Y.S. Proteome: T.Z. and R.S. Organoids: T.Z. and T.J. Visualization: T.Z., and S.Y.S. Supervision: L.K.N. Manuscript writing and editing: T.Z., S.Y.S and L.K.N. Funding: L.K.N.

## Acknowledgements

The authors thank Dr. Oleks Chernyavskiy, Dr. Jihane Homman-Ludiye and Dr. Stephen Firth from Monash Micro Imaging for providing microscopy and image analysis support, Joel Steele, Anup Shah and Iresha Hanchapola from Monash Proteomics & Metabolomics Platform for providing proteomic technical support, and Dr. Lee Wong for helpful comments and discussion of the manuscript.

L.K.N was supported by a Research Fellowship from Victorian Cancer Agency (MCRF18026). L.K.N’s laboratory was supported for this work by an Investigator Initiated Research Scheme grant from National Breast Cancer Foundation (IIRS-20–094) and a Venture Grant from Cancer Council Victoria.

