## Supplementary Information for "Ribosomal Biogenesis Hyperactivation and ErbB signalling Mediated Network Rewiring Causes Adaptive Resistance to FGFR2 Inhibition"

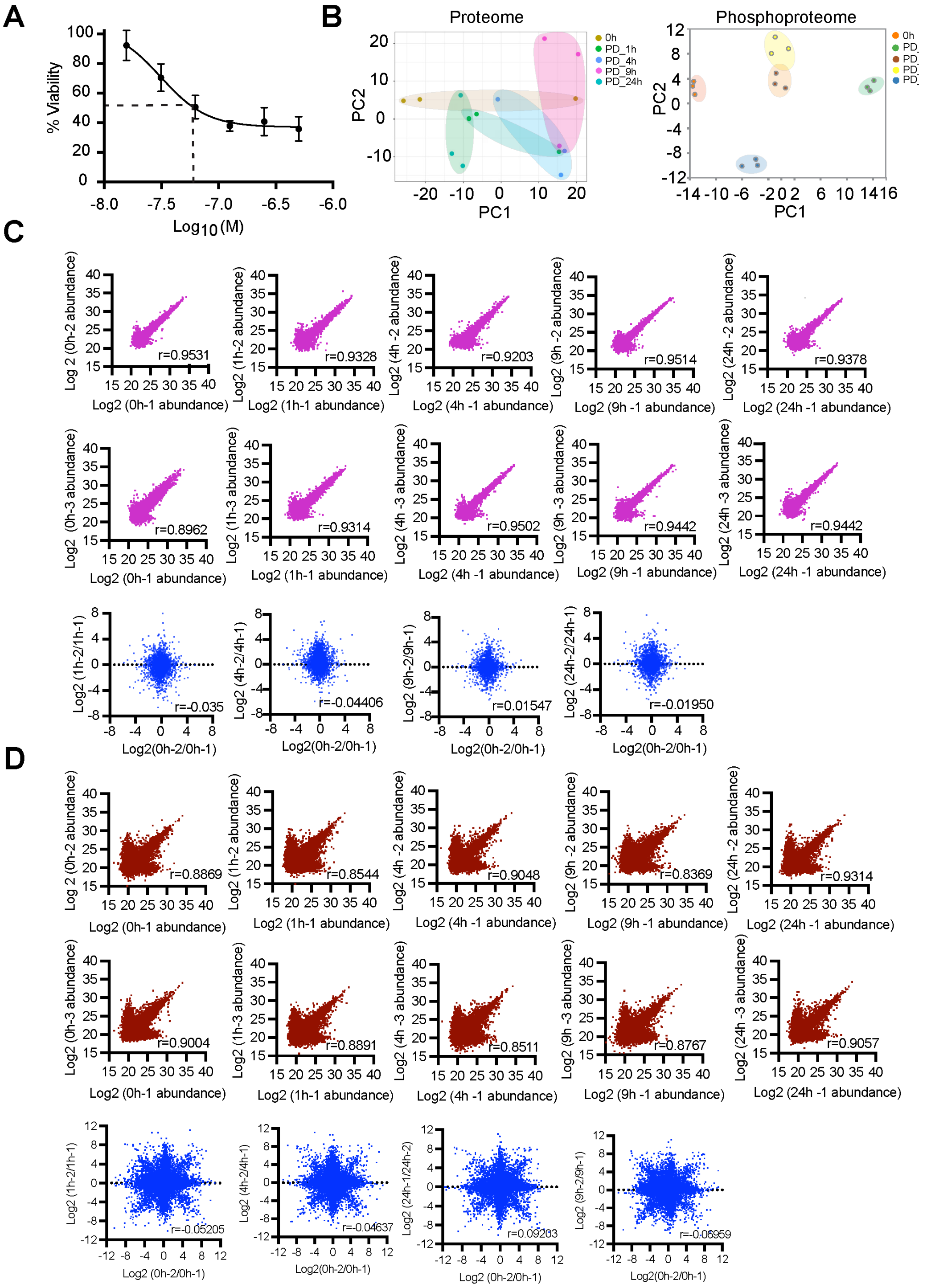


**Figure S1. Quality control of proteomic and phospho-proteomic datasets. (A)** IC50 of PD173074 against MFM223 cells determined using cell viability assays at 96 hours post-treatment. **(B)** Principal component analysis of proteome and phosphoproteome data. Proteome and phosphoproteome datasets showed tight clustering of biological replicates. **(C-D)** Correlation analysis of biological replicates and alterations withing proteome (C) and phosphoproteome datasets (D). High correlation (r > 0.9) was observed between biological replicates, while no correlation (r < 0.05) was detected between different time points, validating the reliability of the profiled dataset.

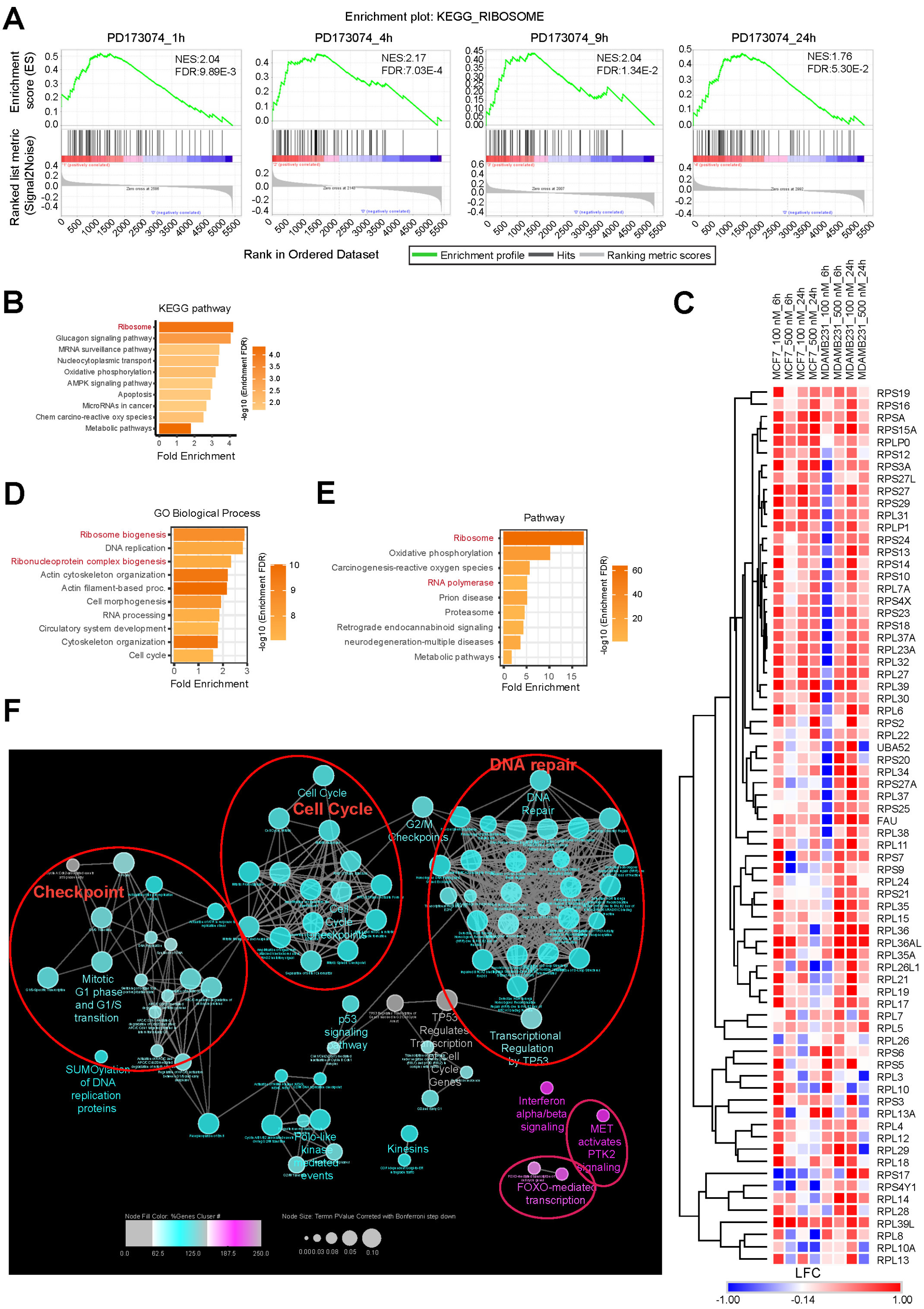


**Figure S2. Upregulation of ribosome biogenesis in response to FGFR inhibition. (A)** GSEA analysis revealed a highly significant enrichment of KEGG ribosome pathway in MFM223 cells treated with PD173074 across all timepoints examined. NES indicates normalised enrichment score, FDR indicates false discovery rate. **(B)** Upregulation of ribosome large and small subunits in MCF-7 and MDA-MB-231 cells treated with PD173074. mRNA expression data were extracted from the Connectivity Map platform (CMAP). **(C)** Gene ontology analysis of 1529 upregulated differentially expressed genes (DEGs) in PD173074-

resistant RT112 cells compared to PD173074-treated parental RT112 cells (24 h) revealed significant enrichment of ribosome biogenesis (p-value < 0.01, LFC > |1|). Datasets were extracted from GSE201395. **(D)** KEGG pathway analysis of 820 up-regulated DEGs in futibatinib -treated brain tumour glioblastoma (BTG) patient-derived xenograft (PDX) mice revealed significant enrichment of the ribosome pathway (adjusted p -value < 0.05, LFC > 1). Datasets were extracted from GSE245624. **(E)** Functional clustering of differentially expressed proteins in KARS inhibitor-treated MiaPaCa-2 cells revealed no ribosome biogenesis-related modules, unlike PD173074-treated MFM223 cells.

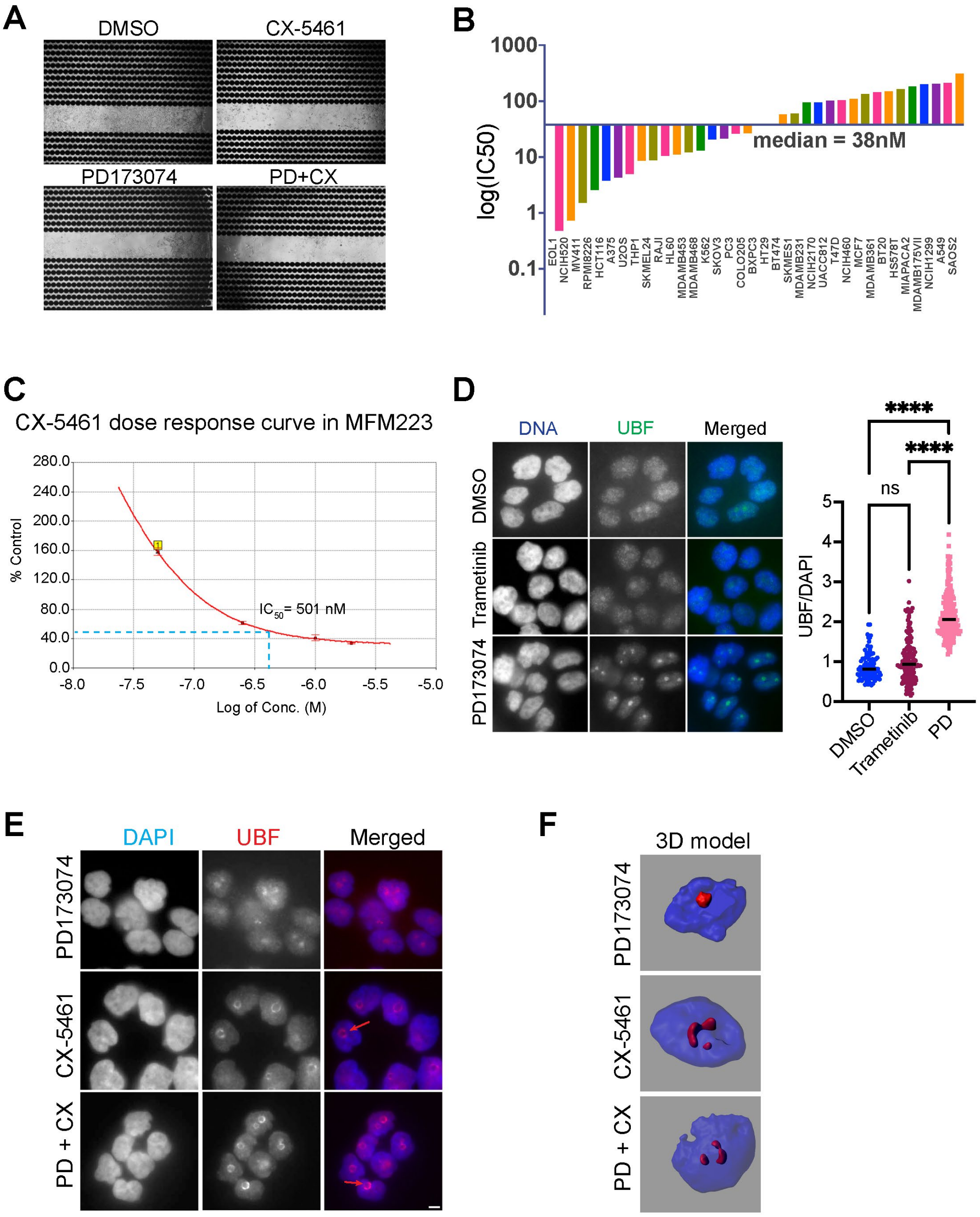


**Figure S3. Upregulation of ribosomal DNA transcription by PD173074. (A)** E-plates used in Figure 4D were imaged after 120 hours (end of the experiment). Images show MFM223 cells treated with 100 nM PD173074, 1 µM CX-5461, 100 nM PD173074 + 1 µM CX-5461 and DMSO. **(B)** The median IC50 values of CX-5461 in various cancer cell lines was 38 nM. **(C)** CX-5461 exhibited a relatively higher IC50 value (501 nM) in MFM223 cells, as determined by xCELLigence RTCA cell proliferation assays. **(D)** Immunofluorescence staining with UBF, with DAPI staining for DNA contents. MFM223 cells were treated with 20 nM PD173074 or 500 nM Trametinib for 4 hours, with DMSO as controls. Representative images of three independent experiments (left panel). Quantification of UBF intensity relative to DAPI intensity (right panel). **** p-value $<0.001$, and ns indicates no significance. Error bars represent mean ± standard error of three independent experiments. At least 150 cells were measured in each three independent experiments. **(E-F)** Immunofluorescence staining with UBF, with DAPI staining for DNA contents. MFM223 cells were harvested as described in Figure 5A. After pre-permineralization, cells were stained with UBF (E). Images were deconvoluted, and 3D reconstructed models are shown (F).

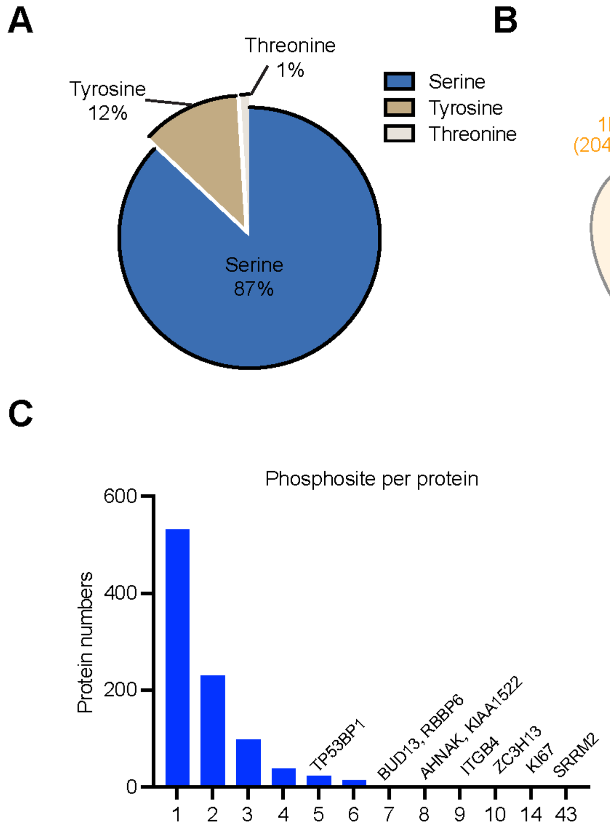


**Figure S4. Dynamic and diverse cellular changes of Phosphoproteomic profiling. (A)** Amino acid composition at the identified phosphosites. Serine phosphorylation predominates, followed by tyrosine and threonine. **(B)** Dynamic change of phosphosites over times. **(C)** Distribution of phosphosites per protein. A subset of proteins, such as TP53BP1, BUD13, and RBBP6, exhibit exceptionally high numbers of phosphosites.

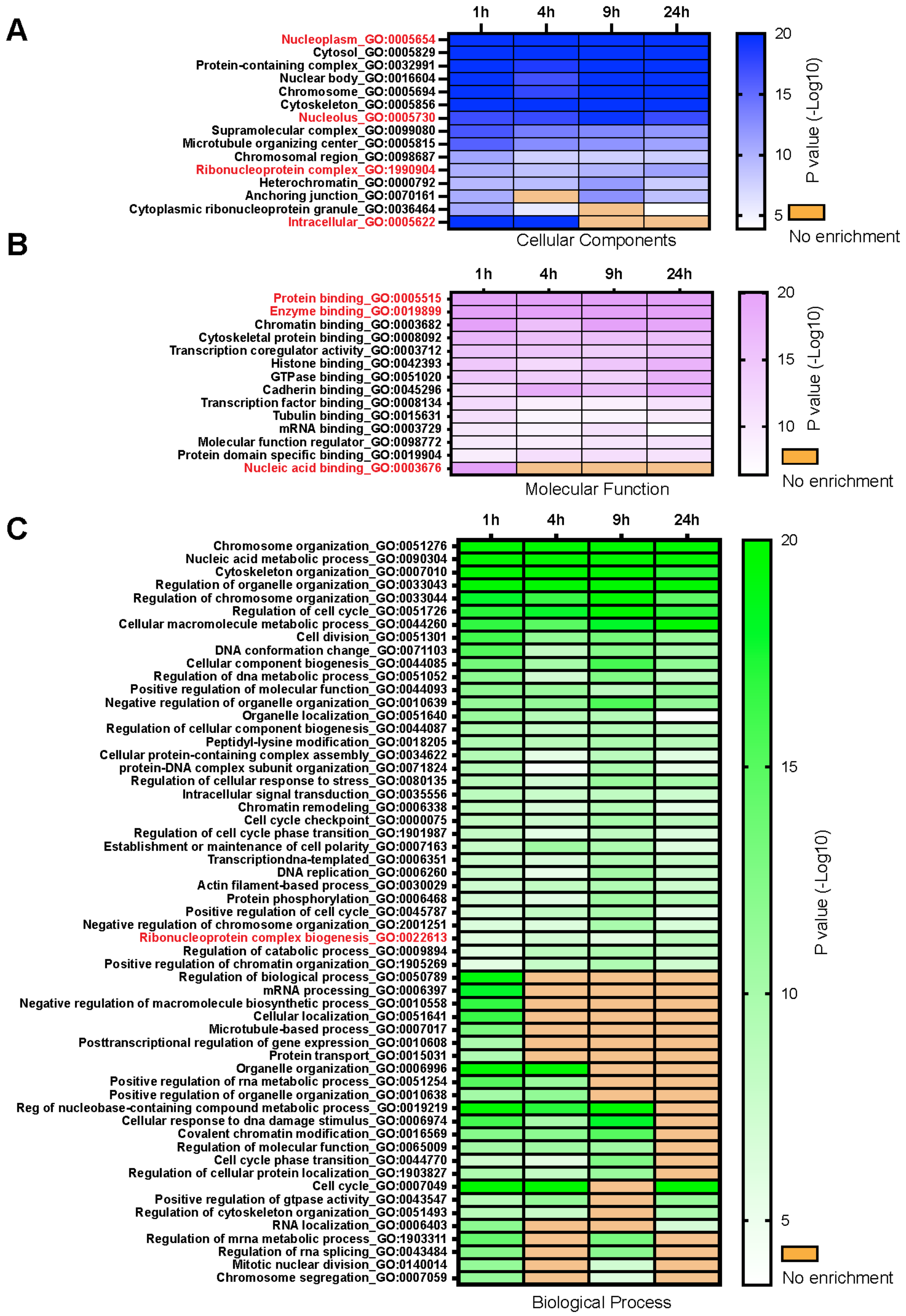


**Figure S5. Gene Ontology analysis of the phosphoproteome profile. (A)** Cellular components. **(B)** Molecular function. **(C)** Biological process. The enrichment terms were displayed for p value (-Log10) > 10 at any timepoint.

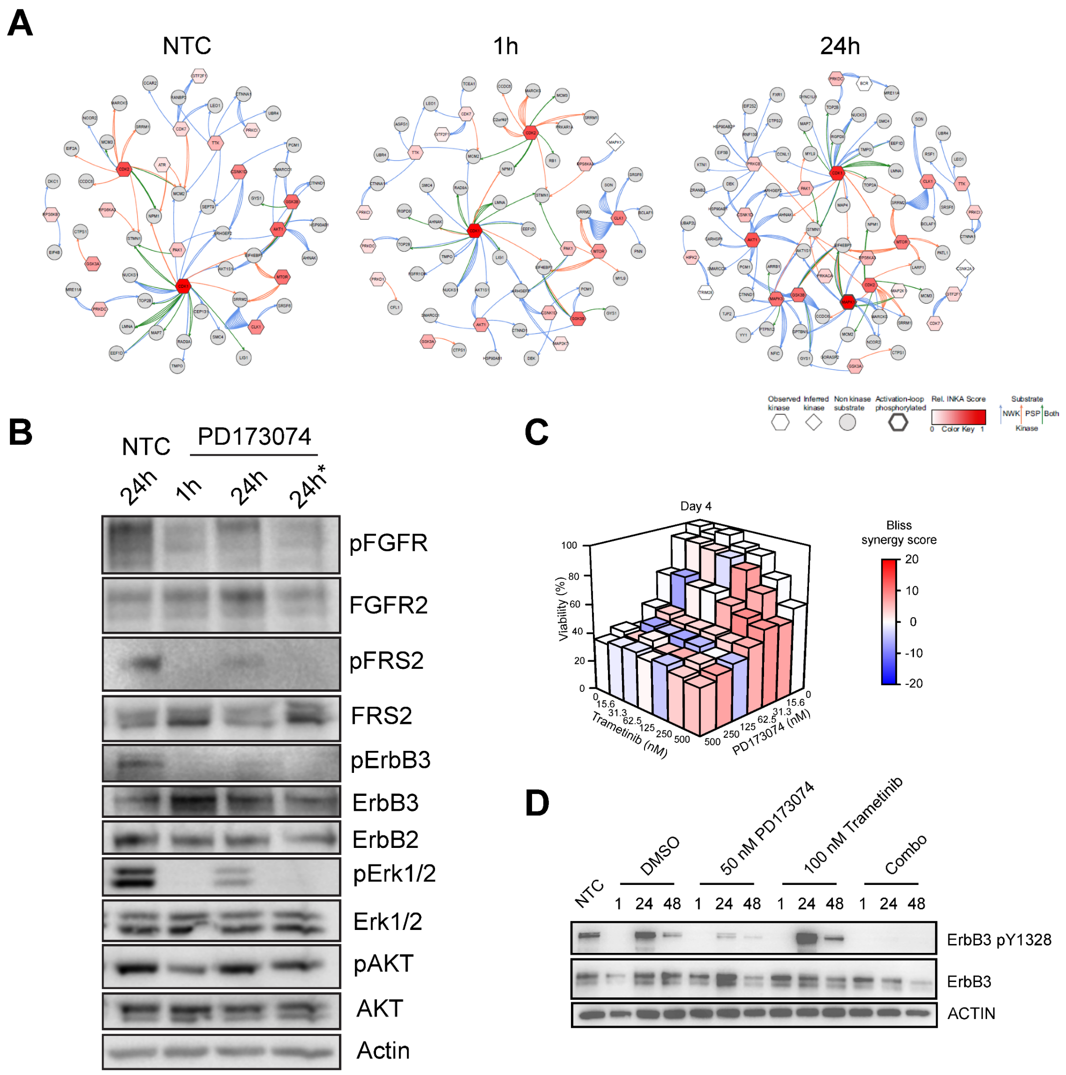


**Figure S6. Reactivation of ErbB/MAPK pathways. (A)** Kinase and substrates network analysis. Reactivation of kinases AKT1, MAPK1, MAPK3 and GSK3B was highlighted in MFM223 cells following PD173074 treatment. **(B)** Reactivation of ErbB and ERK signalling. MFM223 cells were treated with 20 nM PD173074 for 24 hours. Supernatants were collected and replenished into fresh untreated cells for an additional 24-hour treatment (24h*). **(C)** Cell viability and synergy scores quantified for combined treatments of PD173074 and Trametinib. Cell viability from MTS assays were determined at 4 days post-treatment of PD173074 (0 - 500 nM) and Trametinib (0 - 500 nM). The synergy score was calculated based on Bliss' independent model. **(D)** Synergistic effect of PD173074 and Trametinib on pErbB3. MFM223 cells were treated with 50 nM PD173074, 100 nM Trametinib, or their combination (Combo). The ErbB3 phosphorylation was measured at 1 h, 24 h and 48 h post-treatment, with NTC and DMSO treatment as controls. Representative of three independent experiments.

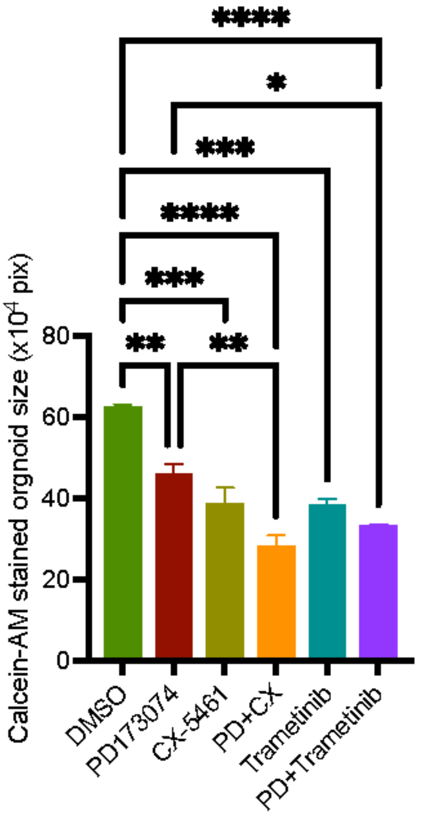


**Figure S7. Effect of combinational treatment on TNBC patient-derived organoids.** Organoid size (×$10^4$ pixels), quantified from Calcein-AM staining (Fig. 7E). ** p-value < 0.01, * p-value < 0.05, and ns indicates no significance. Error bars represent mean ± standard error of three independent experiments.

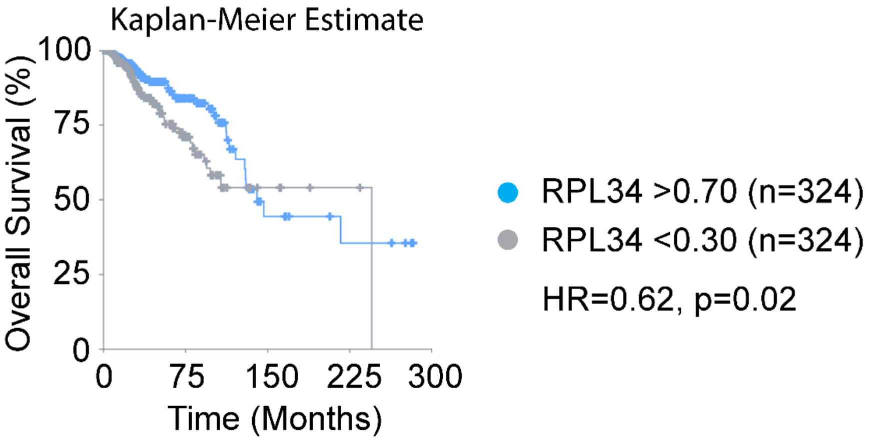


**Figure S8.** Survival analysis of breast invasive carcinoma patients (TCGA, Pan-Cancer Altas). High RPL34 expression was associated with a significantly lower risk of death compared to low expression (p = 0.02, HR = 0.62).

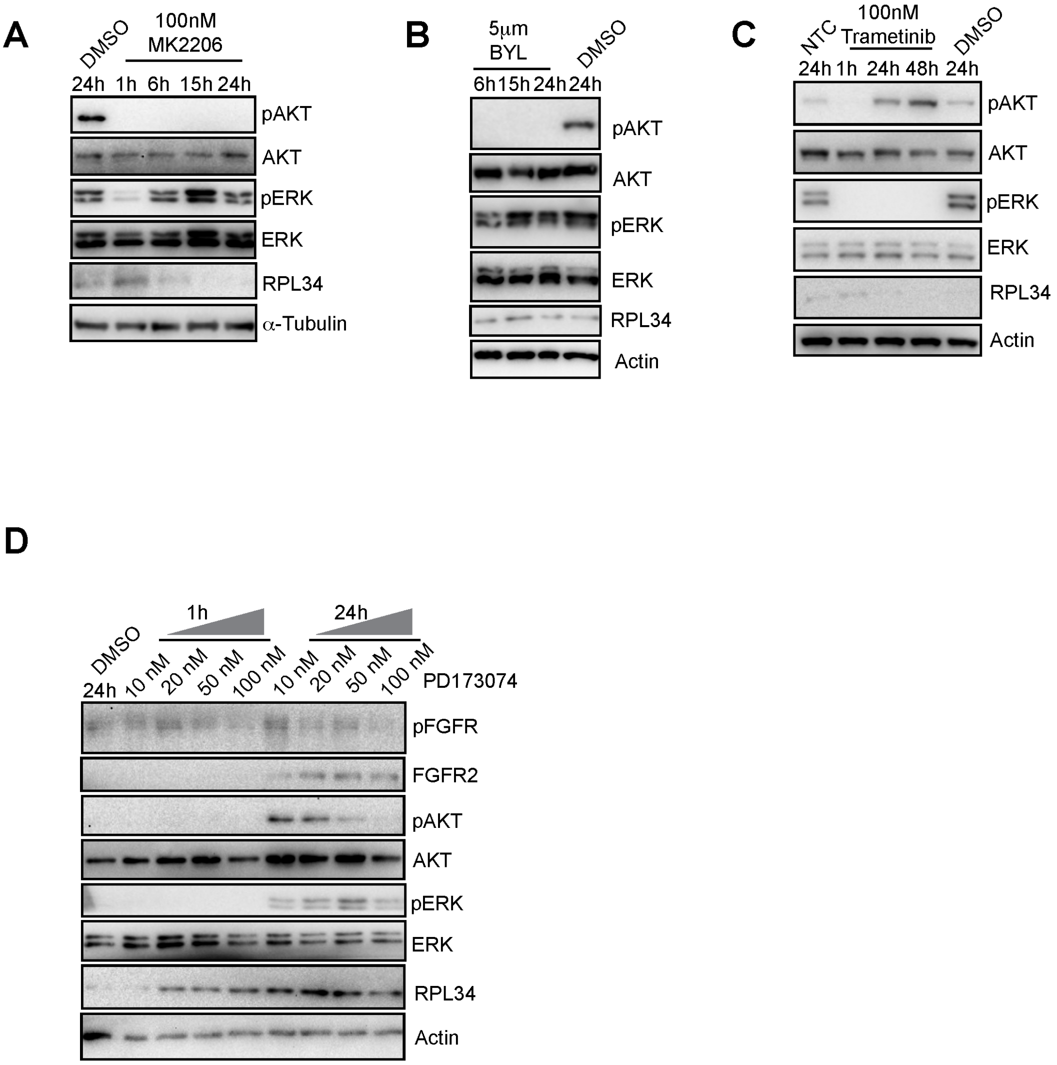


**Figure S9. Crosstalk between the MAPK and AKT pathways. (A)** MEK activation following AKT inhibition. MFM223 cells were treated with 100 nM MK2206, with DMSO as a control. Cells were harvested at 1, 6, 15, and 24 hours post-treatment. Representative of three independent experiments. **(B)** Reactivation of ERK following PI3K inhibition. MFM223 cells were treated with 5 μM BYL719, with DMSO as a control. Cells were harvested at 6, 15, and 24 hours post-treatment. Representative of three independent experiments. **(C)** AKT activation following MEK inhibition. MFM223 cells were treated with 100 nM Tramentinb, with DMSO as a control. Cells were harvested at 1, 6, 15, and 24 hours post-treatment. Representative of three independent experiments. **(D)** Reactivation of AKT and ERK and upregulation of FGFR2 and RPL34. MFM223 cells were treated with 10 nM, 20 nM, 50 nM, 100 nM PD173074 for 1 hour and 24 hours, with DMSO as a control. Representative of three independent experiments.

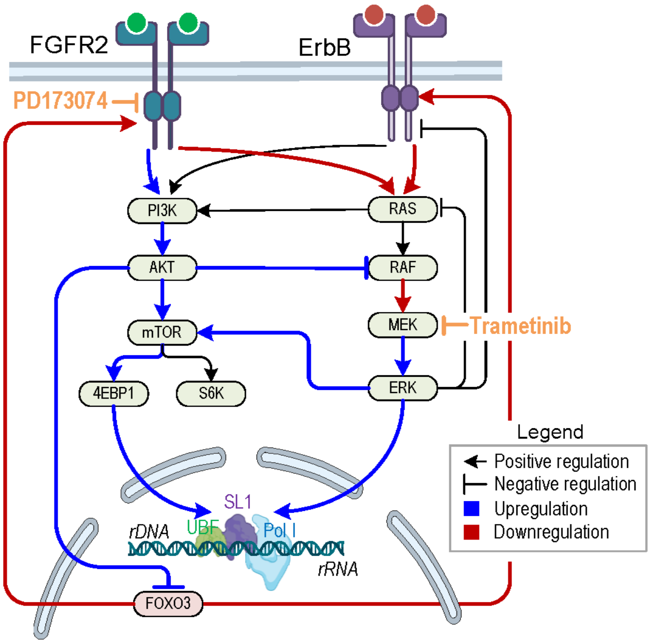


**Figure S10. MEK inhibition blocks the resistance to FGFR2 inhibition.**
MEK inhibition suppress ribosomal biogenesis by blocking AKT-mediated crosstalk and subsequent transactivation of MEK.

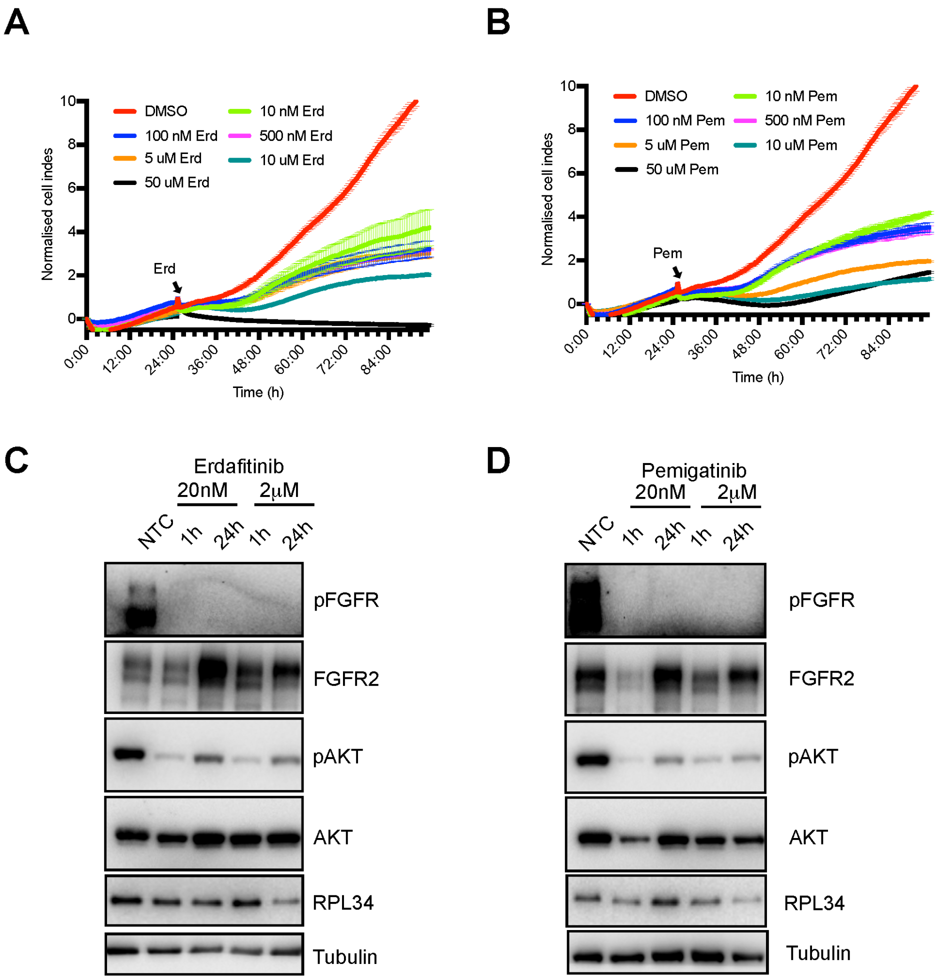


**Figure S11. Effect of newer FGFR inhibitor Erdafitinib and Pemigatinib on MFM223 cells.** Cell proliferation of MFM223 cells treated with 10 nM, 100 nM, 500 nM, 5 μM, 10 μM and 50 μM Erdafitinib **(A)** or Pemigatinib **(B)** for 72 hours, with DMSO as controls, measured with xCELLigence RTCA system. Error bars represent mean ± standard error of three independent experiments. Cell index was normalised to the time when the treatment occurred (indicated by an arrow). **(C-D)** Reactivation of AKT and upregulation of FGFR2 by newer FGFR inhibitors. MFM223 cells were treated with 20 nM or 2 μM Erdafitinib (C) or Pemigatinib (D), with NTC as controls. Representative of two independent experiments.
